# Processive translocation of cohesive and non-cohesive cohesin *in vivo*

**DOI:** 10.1101/2021.10.07.463478

**Authors:** Melinda S. Borrie, Marc R. Gartenberg

## Abstract

Cohesin is a central architectural element of chromosomes that regulates numerous DNA-based events. The complex holds sister chromatids together until anaphase onset and organizes individual chromosomal DNAs into loops. Purified cohesin translocates along DNA in a diffusive fashion that can be propelled by transcribing RNA polymerase. The complex also extrudes DNA loops in a process that consumes ATP. In this study we examine processive genomic translocation of cohesin in vivo. To this end, obstacles of increasing size were tethered to DNA to act as roadblocks to complexes mobilized by transcription in yeast. The obstacles were built from a GFP-lacI core fused to one or more mCherries. A chimera with four mCherries blocked cohesin passage in late G1. During M phase, the threshold barrier depended on the state of cohesion: non-cohesive complexes were also blocked by four mCherries whereas cohesive complexes were blocked by as few as three mCherries. Furthermore, cohesive complexes that were stalled at obstacles, in turn, blocked the passage of non-cohesive complexes. That synthetic barriers alter cohesin redistribution demonstrates that the complex translocates processively on chromatin in vivo. The approach provides a relative measure of the maximum size of the DNA binding chamber(s) of cohesin. Together, this study reveals unexplored limitations to cohesin movement on chromosomes.

**Significance Statement:** Cohesin is an architectural protein that brings distant chromosomal DNA sites together. The complex links sister chromatids cohesion but it also binds to single pieces of DNA in ways that do not generate cohesion. One class of non-cohesive complexes organizes chromosomal DNA into loops. All cohesin complexes move on DNA but the constraints on such movement are not fully explored. Here, we use size-calibrated obstacles in yeast to interrogate cohesin and the properties of its movement on DNA. We show that both cohesive and non-cohesive complexes translocate processively on chromosomes. In addition, we show that cohesive and non-cohesive complexes are blocked by obstacles of different size. Lastly, we show that stalled cohesive complexes block passage of non-cohesive complexes.

## Introduction

Cohesin is a central architectural element of chromosomes that functions by bringing distant DNA sites together (1). The protein complex was first characterized for its role in sister chromatid cohesion. Cohesin is now known to participate in most if not all fundamental chromosomal processes, including transcription, replication and repair. Recent work has shown that the complex also organizes chromosomes into large DNA loops that contribute to the segmentation of the genome into self-associating domains. Mis-regulation of cohesin has been identified as the cause of certain developmental diseases known collectively as cohesinopathies (2). Mutations in cohesin subunits have also been associated with human cancers.

The evolutionarily conserved core cohesin complex contains two SMC subunits (Smc1 and Smc3) and an *α* kleisin subunit Mcd1 (also known as Scc1/Rad21). The proteins assemble into a ring that can embrace DNA topologically, most notably when engaged in sister chromatid cohesion (3, 4). In yeast, the complex accumulates densely across tens of kilobases surrounding centromeres (5, 6). Along chromosome arms, the complex binds between convergently transcribed genes. Chromosome conformation capture (3C) techniques have shown that such sites participate both in sister chromatid cohesion and chromosomal loop formation in yeast (7–10). Cohesin performs these same architectural functions in higher eukaryotes (11–13).

Cohesin loading onto yeast chromosomes by NIPBL (Scc2 in yeast) begins each cell cycle in late G1. Initially, the interaction is dynamic. The complex cycles on and off DNA because loading is balanced by an unloading factor, WAPL (Wpl1 or Rad61 in yeast). During S phase, Eco1 (also known as Ctf7) halts cohesin loss by acetylating the Smc3 cohesin subunit, which antagonizes Wpl1 (14–18). Eco1 travels with the DNA replication fork, presumably to couple the stabilization of cohesive complexes directly with the creation of sister chromatids (19).

A second dynamic property of cohesin is the ability to redistribute on chromosomes. Cohesin accumulates at genomic sites that are often great distances from where the complex initially loads (20). Initially, RNA polymerase was seen as a central driver of repositioning because cohesin on dormant or weakly transcribed genes was relocated downstream by transcriptional induction (5, 6, 21). At least three observations suggested that cohesin relocates by processive translocation: 1) at low temperature, gene induction moved the complex to intermediate positions between initial and final binding sites (22); 2) cohesin arriving at new sites following changes in transcription did not require the Scc2 loader for accumulation (22, 23); 3) cohesin mobilized by an ectopic polymerase in yeast redistributed to the end of the DNA travelled by the polymerase (24). Indeed, cohesin was shown to diffuse along DNA *in vitro*, and transcribing RNA polymerase was shown to bias the directionality of movement (24–26). More recent biochemical work showed that cohesin, in combination with NIPBL/Scc2, possesses an intrinsic motor activity that enables translocation of the complex along DNA in an ATP-dependent fashion (27, 28). Importantly, these studies showed that translocating cohesin can extrude DNA loops, an activity that is thought to form the large chromosomal loops that typify genomes (29–31). In support of the loop extrusion model, the size and distribution of chromosomal loops are altered by factors that influence the residence time of cohesin on chromatin (8, 9, 32–34).

We recently obtained evidence for processive cohesin translocation on chromosomal DNA in yeast using a system involving transcriptional induction of a cohesin bound gene (35). Upon induction of the gene, cohesion of the locus was lost. If, however, the locus was first converted to an extrachromosomal circle, cohesion persisted. The results are most readily explained by redistribution of topologically-bound cohesin complexes by processive translocation because such complexes cannot migrate off circularized DNA. In the present study, the properties of transcription-propelled complexes were further studied by measuring their ability to bypass synthetic obstacles of increasing molecular size. This work provides direct evidence that cohesin translocates processively on DNA *in vivo*. Moreover, the data reveal that the fate of encounters with barriers depends on the cohesive state of the complex: a barrier sufficient to trap cohesive complexes is not large enough to trap non-cohesive complexes.

## Results

### Nuclear membrane tethering of DNA blocks cohesin translocation

Our previous study found that DNA circularization, as well as downstream convergently oriented genes, prevented transcription-driven cohesion loss (35). As stated above, we inferred that circles trapped cohesin that was continuously mobile. Downstream convergent genes, on the other hand, were seen simply as barriers to cohesin passage. Intriguingly, DNA-bound lac repressors did not block escape of cohesin (Figure 1A). We hypothesized that the bacterial protein (lacI fused to GFP) was smaller than the chamber of the complex through which DNA passes. If topologically-bound cohesin truly redistributes by translocation *in vivo*, then DNA-bound proteins of sufficient size should block passage of the complex. To begin to explore this concept, we used a GFP-lacI chimera linked to FFAT, a tetrapeptide motif that associates with the inner nuclear membrane (36). When simultaneously tethered to DNA and the nuclear membrane, GFP-FFAT-lacI represents an immobile obstacle of essentially infinite size (Figure 1A).

**Figure 1.**
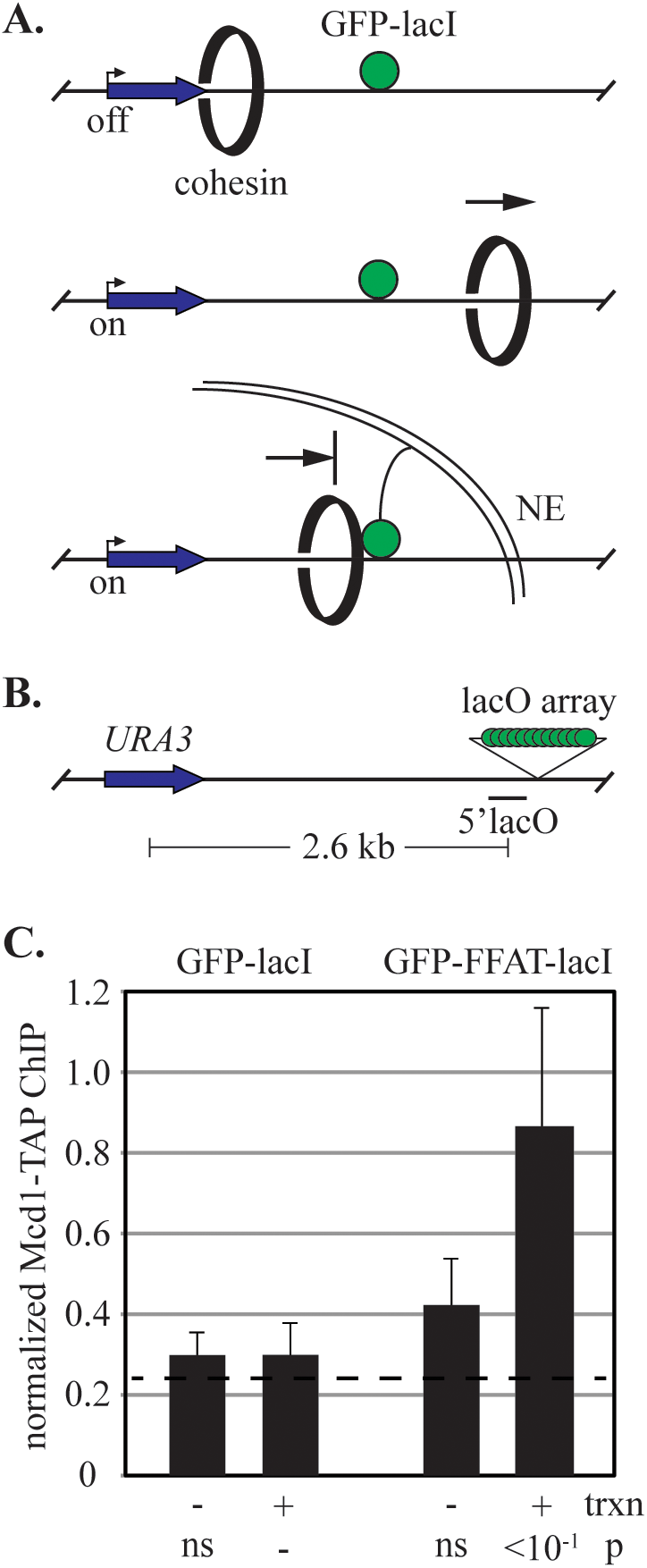
Anchoring DNA to the inner nuclear membrane blocks cohesin translocation. **A)** Schematic of transcription-triggered cohesin translocation and blockage by membrane anchorage. Only one GFP-lacI is shown for clarity. NE = nuclear envelope. **B)** Schematic of the locus used to test translocation. Binding of Mcd1 was evaluated at the 5’lacO site immediately upstream of the lacO array. **C)** ChIP-qPCR of Mcd1-TAP. Strains MSB114 and MRG6947 were arrested in M phase by Cdc20 depletion and then *URA3* transcription was induced. Values were normalized to a cohesin bound site (see Materials and Methods). Measurements at a cohesin-free site (534 kb on Chr IV) were averaged for all experiments of the figure panel and shown as a dashed line. p values for pairwise Student’s t tests are presented relative to benchmark sample (designated with a dash). ns, not significant.

Translocation assays were performed at an engineered genomic locus (Figure 1B) that contains a 256 lac operator array downstream of *URA3*, a gene that can be induced 3-5 fold by withholding uracil. Previous work showed that cohesive complexes associate with uninduced *URA3* until transcriptional induction triggers their departure. Cohesin levels at the gene do not diminish, however, because new complexes reload during the experimental interval ((35), see example in figure S2). Here, chromatin immunoprecipitation coupled with quantitative real-time PCR (ChIP-qPCR) was used to monitor arrival of cohesin at a downstream site immediately preceding the lac operator array (hereafter referred to as the 5’lacO site). Immunoprecipitations were performed with antibodies against a TAP epitope affixed to Mcd1.

*URA3* transcription was induced after cells were arrested in M phase by depletion of Cdc20. When GFP-lacI was expressed, cohesin at 5’lacO was low both before and after *URA3* induction (Figure 1C). The result is consistent with the inability of GFP-lacI to block translocating cohesin complexes. Expression of GFP-FFAT-lacI, on the other hand, increased the level of cohesin at 5’lacO following transcription. This finding serves as a proof-of-principle that large immobile objects block translocation of cohesin complexes in yeast. In *B. subtilis*, tethering a DNA-bound repressor to the plasma membrane similarly blocked translocation of bacterial SMC proteins (37).

### Large DNA-bound proteins block cohesin translocation

To determine if large proteins could similarly block cohesin passage, a series of size-calibrated obstacles was generated by fusing increasing numbers of mCherry to GFP-lacI (Figure 2A-C). This red fluorescent protein is ideally suited as a size standard because it has a compact monomeric structure unlike its tetrameric progenitor, DsRed (38–40). Like GFP-lacI, the mCherry chimeras bind lacO arrays, albeit with reduced efficiency, as demonstrated by weaker GFP foci and higher background nuclear fluorescence (Figure S1).

**Figure 2.**
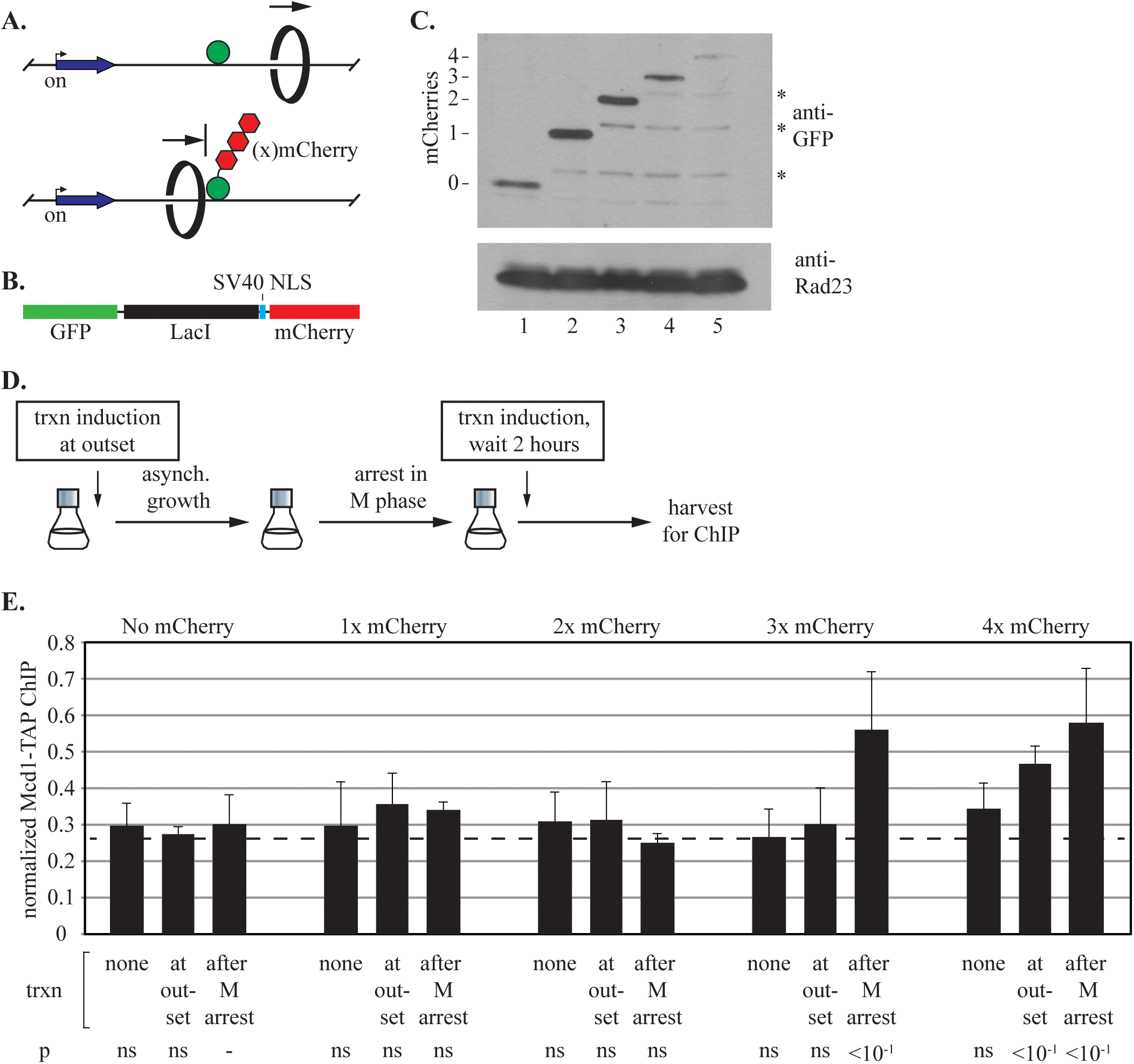
Synthetic barriers of sufficient size block cohesin translocation. **A**) Schematic of transcription-triggered cohesin translocation and blockage by GFP-lacI chimeras. **B**) Organization of the GFP-lacI chimeras with mCherries. **C**) Immunoblot of GFP-lacI chimeras. Strains expressing MSB174, MSB179, MSB180, MSB181 and MRG7102 were used. The lower molecular weight bands marked with * likely represent partial proteolytic cleavage of mCherry (72). **D**) Experimental flow chart. Transcription was either induced from the outset of growth, after M phase arrest or not at all. **E**) ChIP-qPCR of Mcd1-TAP. Cohesin binding at the 5’lacO site was evaluated in strains MSB114, MRG6810, MRG6812, MSB165.1 and MRG7060.

Accumulation of cohesin at 5’lacO was measured in M phase-arrested cells following three different *URA3* induction protocols (Figure 2D). In the first, *URA3* was never induced. Accordingly, cohesin levels at 5’lacO remained low and comparable to the cohesin-free control site, which is represented by a dashed line in the figures. In the second protocol, *URA3* was induced following imposition of an M phase arrest. In this case, accumulation of cohesin at 5’lacO depended on the identity of the GFP-lacI chimera expressed (Figure 2C). When GFP-lacI was fused to one or two mCherries or none at all, cohesin levels did not rise above the baseline. By contrast, when GFP-lacI was fused to three or four mCherries cohesin accumulated at 5’lacO. A more thorough analysis of the cohesin distribution around the engineered locus is shown in Figure S2. In the three mCherry strain, the most dramatic changes in cohesin binding occurred only between *URA3* and the lacO array, as expected for complexes travelling from the induced gene into a downstream roadblock. No dramatic changes were found with GFP-lacI alone, including at sites downstream of the array. This suggests that cohesin distributes throughout the region in the absence of a single strong barrier. To be certain that an aberrant property of mCherry was not responsible for 5’lacO accumulation, the experiment was repeated with a chimera bearing three copies of monomeric yeGFP. Similar results were obtained (Figure S3). These results indicate that DNA bound proteins of sufficient size block passage of translocating cohesin complexes.

In the final experimental protocol, *URA3* was induced at the outset of asynchronous growth. Previously we found that a downstream convergently-oriented gene could not block translocation of cohesive complexes under these conditions. The basis for this phenomenon was not understood (35). When *URA3* was induced from the outset here, most of the barrier chimeras similarly failed to accumulate cohesin at 5’lacO. Some binding, however, was detected when GFP-lacI was fused to four mCherries (Figure 2E). The data suggests that different sized barriers act as obstacles under different conditions. Additional experiments elaborate on this concept below.

### Trapped cohesin complexes remain cohesive

To determine whether the complexes at 5’lacO hold sister chromatids together, we evaluated cohesion of the chromosomal domain. In our strains, target sites for the R site-specific recombinase flank the entire region (Figure 3A). Induction of the recombinase during an M phase arrest generates a pair of DNA circles, one from each sister chromatid. Owing to the bound GFP-lacI chimeras, a single dot of fluorescence appears if the DNA circles are cohesed whereas a pair of dots appears if they are not (Figure S1). *URA3* transcription was induced according to the three protocols described above and cohesion was evaluated at the end of the growth protocol (Figure 3B). In the absence of induction, cohesion of DNA circles was observed in all strains due to the presence of cohesive cohesin on *URA3* (Figure 3C; (35)). When transcription was induced following M phase arrest, cohesion was greatly reduced in strains with chimeras bearing one or two mCherries or none at all. This result indicates that these chimeras are too small to block translocating cohesin. By contrast, cohesion persisted when transcription was induced in strains expressing chimeras with three and four mCherries. Thus, consistent with the results in figure 2, DNA-bound chimeras equal to or larger than three mCherries block passage of translocating cohesive cohesin. We conclude that at least some of the trapped cohesin complexes detected by ChIP-qPCR are engaged in sister chromatid cohesion.

**Figure 3.**
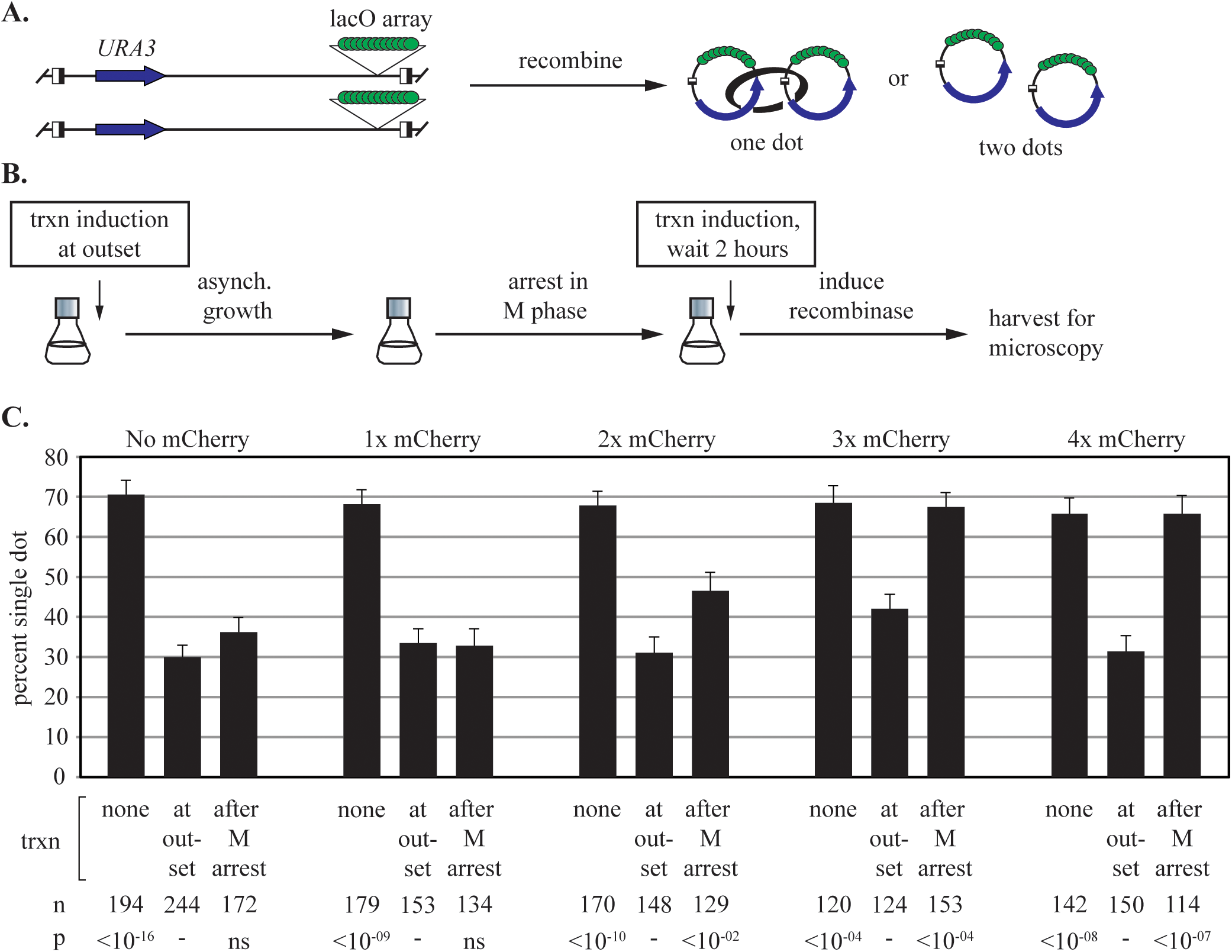
Synthetic barriers of sufficient size block translocation of cohesive cohesin. **A**) An assay for cohesion at a specific chromosomal location. The GFP-tagged region of interest is converted to extrachromosomal circles by site specific recombination. Half-filled boxes represent target sites for the inducible R recombinase. **B)** Experimental flow chart. **C**) Cohesion analysis of extrachromosomal circles. Strains MSB35.2, MRG6806, MRG6808, MSB167 and MRG7060 were used. p values for pairwise χ2 tests are presented relative to a benchmark condition for each strain (designated with a dash). Error bars represent the standard error of proportion. n = the number of cells examined.

When transcription was induced from the outset of cell growth, cohesion of the DNA circles was abolished in all strains, irrespective of the chimera expressed. This result recapitulates the earlier observation that convergent genes downstream of *URA3* also do not trap cohesive complexes under identical induction conditions (35). These data suggest that the ability of an obstacle to block passage of cohesin depends on the interplay between transcription and cell cycle stage. Data supporting this view is presented below.

### A larger barrier is required to block cohesin translocation in late G1 phase

The studies above examined translocation of cohesin on newly replicated chromatids in M phase-arrested cells. To study translocation on unreplicated chromatin, cells were arrested in late G1 by expressing a non-degradable allele of Sic1, a Cdk1 inhibitor that prevents S phase entry (41). By definition, the chromatin-bound complexes at this stage of the cell cycle are not cohesive. The steps of the growth protocol are depicted in figure 4A and validation of the growth arrest is shown in Figure S4. In this scenario, transcriptional induction caused accumulation of cohesin at 5’lacO only in a strain expressing the four mCherry barrier. In strains expressing three mCherries or GFP-lacI alone, cohesin accumulation matched the background levels of the uninduced controls (Figure 4B). This data suggest that different size barriers are required to block cohesin at different stages of the cell cycle. Apparently, something happens during S phase that affects the how cohesin translocates past obstacles on DNA.

**Figure 4.**
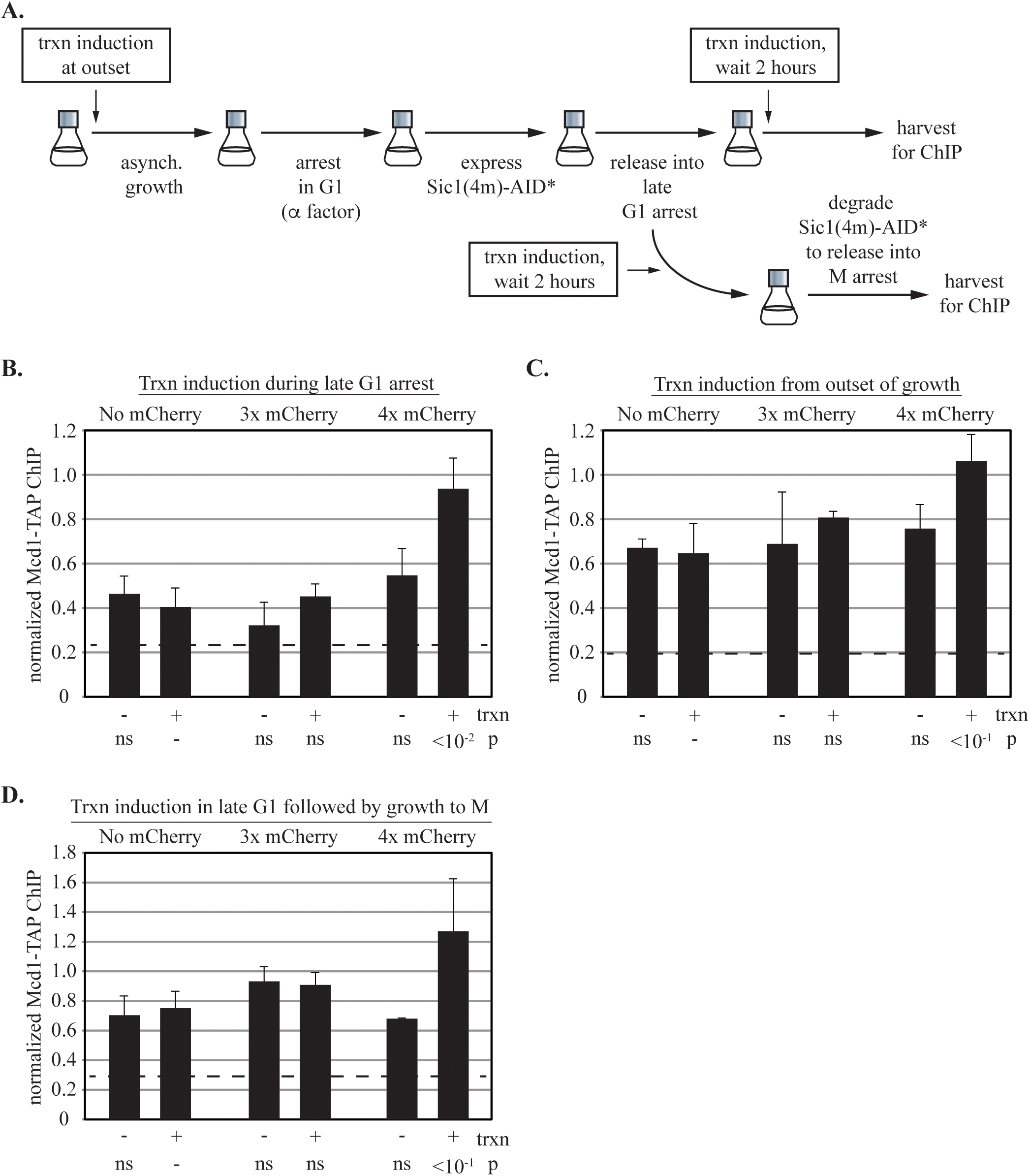
Synthetic barriers to cohesin translocation before and after S phase. **A**) Experimental flow chart. **B**) Barriers to translocation in late G1-arrested cells. ChIP-qPCR of Mcd1-TAP was performed in strains MRG7027, MRG7026 and MRG7059 that were arrested by expression of non-degradable Sic1. *URA3* transcription was induced after cell cycle arrest. **C**) Same as **B** except transcription was induced from the outset of growth. Two trials were performed for each strain. **D**) Barriers to translocation in cells progressing from G1 to M phase. Details of the cell cycle arrest, release and re-arrest are described in Materials and Methods. Strains MRG7033, MRG7032 and MRG7282 were used.

We hypothesized that the four mCherry threshold seen in figure 4B was related to the same threshold seen during M phase arrest after induction from the outset of growth (Figure 2E). To understand this line of reasoning, recall that cohesin binding resets after each anaphase passage when separase activity subsides and Mcd1 synthesis resumes (42–44). If *URA3* is already transcribing when cohesin binding resumes in late G1 then the ensuing translocation of the complex would occur on an unreplicated chromosome, at least until S phase. Put simply, translocation before DNA replication would bypass the three mCherry limitation normally seen in M phase cells. To test this idea, *URA3* was induced from the outset of growth and barrier activity of the obstacles was analyzed by ChIP-qPCR after cells were arrested in late G1. Figure 4C shows that cohesin accumulated at 5’lacO in cells that expressed a four mCherry obstacle but not the three mCherry variant or GFP-lacI alone. These data indicate that when *URA3* is induced from the outset of growth, transcription triggers translocation of cohesin before DNA replication, leading to mobile complexes that pass obstacles the size of three mCherries but not four mCherries.

### Translocation of complexes that arrive after DNA replication

Cohesin loads continuously onto chromosomes from late G1 to anaphase (19, 35, 45). Complexes that arrive after DNA replication are not cohesive, do not acquire S phase-dependent acetylation and are subject to turnover by Wpl1 (46–50) To gain a more complete picture of cohesin dynamics at *URA3*, transcription was induced while cells were arrested in late G1 and then the cultures were released for growth into a subsequent M phase arrest. ChIP-qPCR at this stage thus reports on those cohesin complexes that load and translocate to 5’lacO before S phase, as well as those that load and translocate after S phase. Formally, this induction protocol also records translocation of complexes that load during DNA replication of the gene, a time span that is likely too short to affect the results significantly. The novel arrest-and-release procedure was achieved by inducible clearance of AID-tagged, “non-degradable Sic1”, using the auxin analog 1-naphthaleneacetic (Figure S4). The ChIP-qPCR data of figure 4D show that when *URA3* transcription was induced, cohesin only accumulated at 5’lacO in the strain expressing the four mCherry obstacle. That the barrier limit did not revert to three mCherries indicates that late arriving, non-cohesive cohesin was not blocked by the smaller barrier.

To examine late arriving complexes exclusively, a TAP-tagged allele of Mcd1 was induced in M phase arrested cells (Figure 5A). Subsequent induction of transcription caused accumulation of TAP-tagged cohesin at the three and four mCherry obstacles (Figure 5B). On the surface, it would appear that late arriving complexes cannot pass the smaller three mCherry barrier. In this iteration of the experiment, however, untagged cohesive cohesin was created, having been loaded earlier in the cell cycle. Accordingly, in this scenario cohesive cohesin stalled at three mCherries might block passage of non-cohesive complexes. To test this notion directly, a second iteration of the late-loading experiment was performed in which pre-existing cohesin was removed during the M phase arrest by auxin-inducible degradation of endogenous Mcd1 (Figure 5A). Depletion of the AID-tagged protein was confirmed in Figure S5. Figure 5C shows that the late arriving TAP-tagged cohesin now bypassed the three mCherry obstacle and was instead blocked by four mCherries (Figure 5C). The accumulation signal was weak but significant. Thus, in the absence of cohesive cohesin, late-arriving, non-cohesive complexes stopped only at the larger barrier to translocation. Importantly, the data show that stalled cohesive complexes block the passage of non-cohesive complexes.

**Figure 5.**
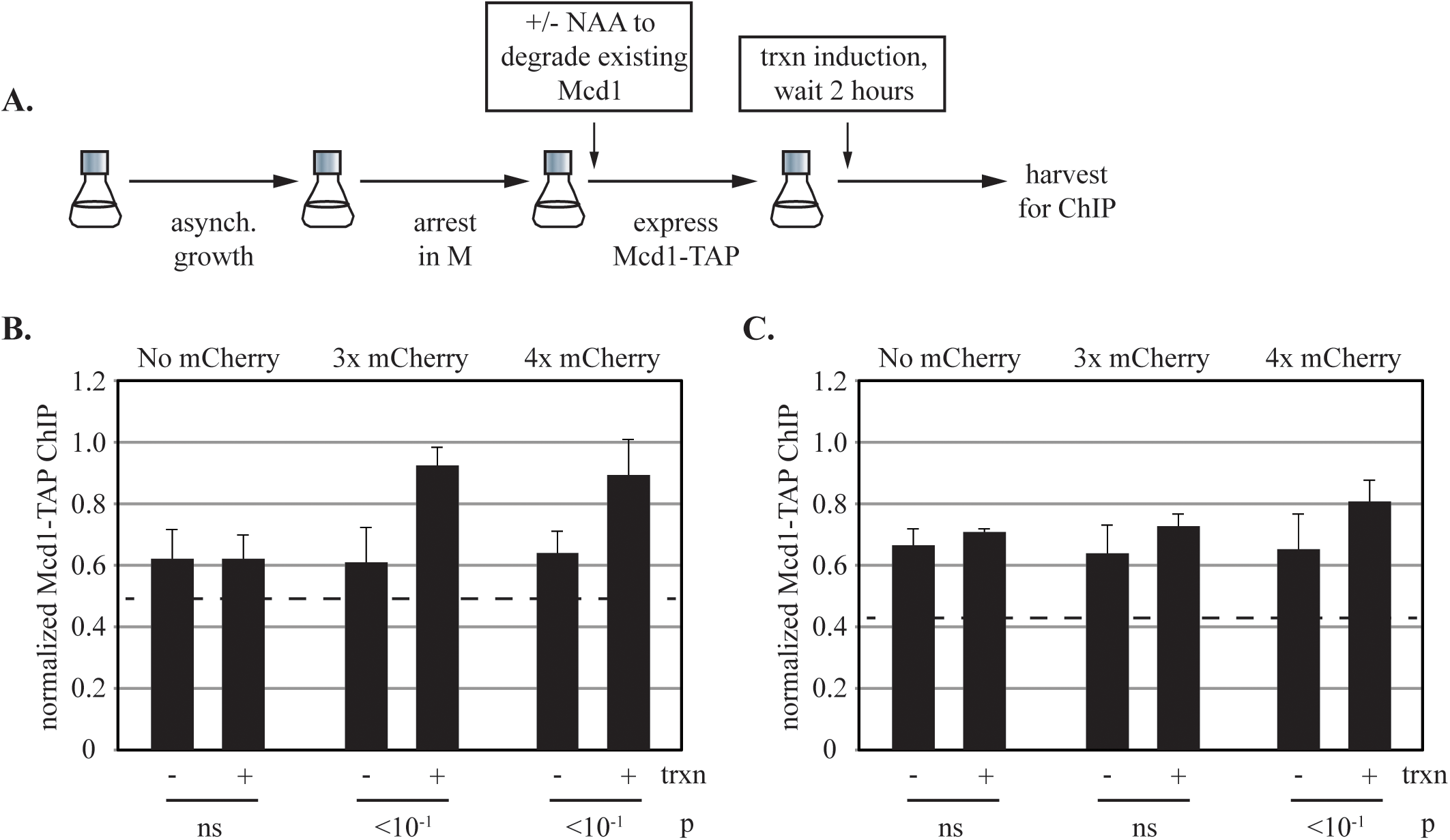
Translocation of cohesin complexes loaded in M phase. **A)** Experimental flow chart. **B)** ChIP-qPCR of Mcd1-TAP. Strains MRG6989, MRG7294, and MRG7292 were used. The Mcd1-TAP allele bears separase cleavage site mutations (R180D,R268D) that should not affect the experimental outcome. The baseline levels of the Mcd1-TAP signal were higher in this experiment relative to figures 2 and 4, likely owing to robust Mcd1 expression from the *GAL1* promoter (45). Nevertheless, the relative levels were internally consistent with the previous figures. **C)** ChIP-qPCR of Mcd1-TAP after depletion of endogenous Mcd1. Strains MRG7317.1, MRG7319.1 and MRG7318.1 were used. See Materials and Methods for timing of each experimental step. Cohesin did not accumulate at the three mCherry obstacle, unlike non- cohesive complexes in figure 5B and cohesive complexes in figures 2E and 3C.

### Eco1 and Wpl1 modulate cohesin’s encounters with barriers

We next tested the influence of regulators of cohesin, Eco1 and Wpl1, on translocation of the complex past synthetic obstacles. The proteins were depleted by auxin inducible degradation during a G1 arrest, then cells were grown to the subsequent M phase arrest before inducing *URA3* (Figures 6A). Clearance of the AID-tagged proteins is shown in figure S5A-C. Transcriptional regulation of *URA3* was unchanged by the depletions (Figures S5D and S6B). In these analyses, ChIP-qPCR data of transcribed and non-transcribed samples were always compared to avoid impact that the mutants might have on absolute levels of cohesin binding.

**Figure 6.**
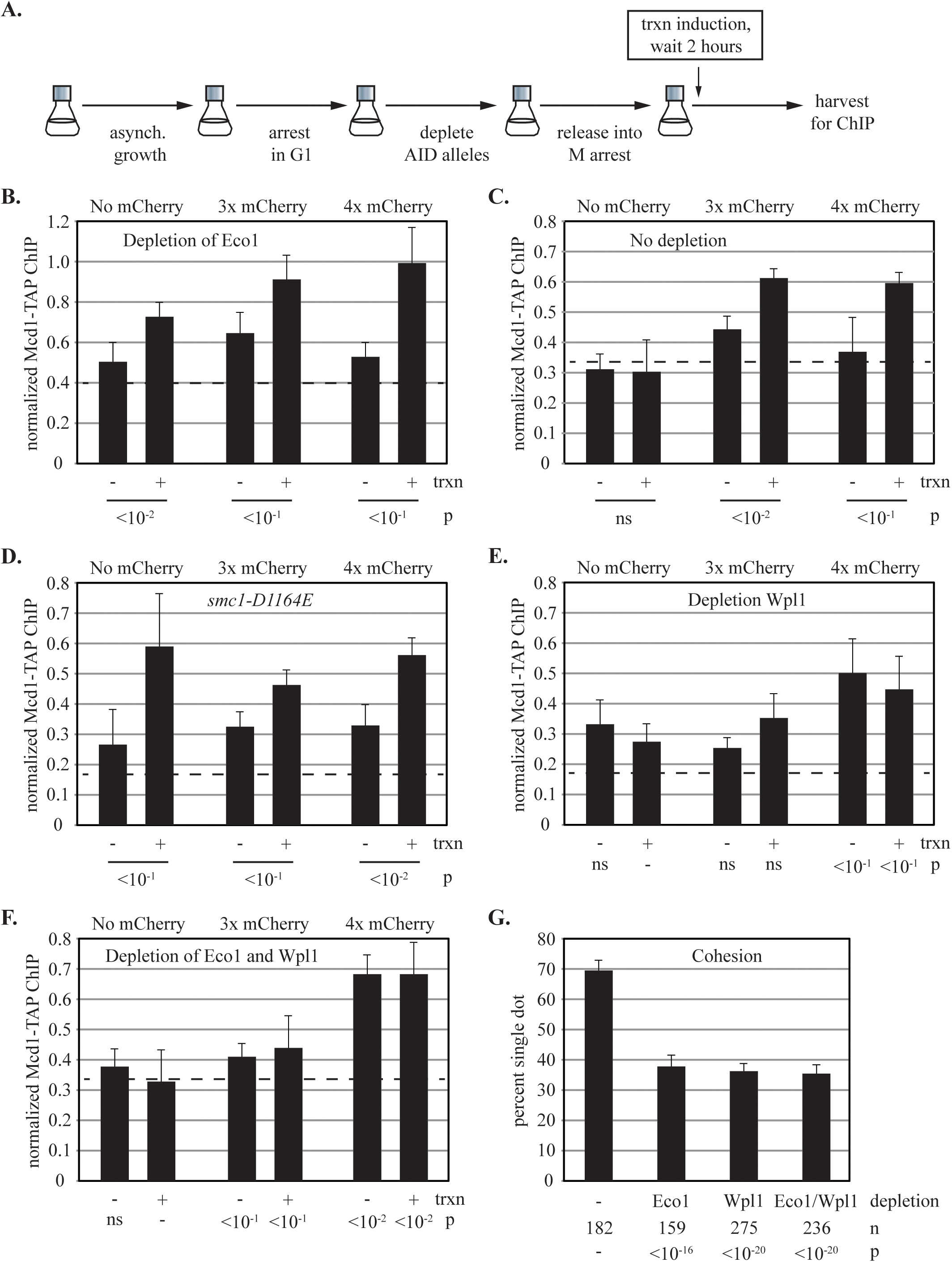
The impact of cohesin regulators on translocation past synthetic barriers. **A**) Experimental flow chart. **B)** Translocation in the absence of Eco1. Eco1-3xmAID was depleted by the addition of 100 µM NAA to strains MRG7301.1, MRG7300.1, MRG7302.1, as described in Materials and Methods and graphically in panel A. p values for pairwise Student’s t-tests are presented for the transcriptional induction of each strain. **C)** Translocation in strains with Eco1. Strains MRG7269.1, MRG7270.1 and MRG7275.1 express OsTir1 but lack AID-tagged genes. **D)** The impact of *smc1-D1164E* on translocation past barriers. Strains MRG7151, MRG7146 and MRG7159 were arrested in M phase before transcription was induced, as in figure 2. **E)** Translocation in the absence of Wpl1. Wpl1 was depleted from strains MRG7327.1, MRG7329.1, MRG7324.1. **F**) Translocation in the absence of Eco1 and Wpl1. Strains MRG7332.1, MRG7356.1 and MRG7353.1 with 3xmAID alleles of *eco1* and *wpl1* were used. **G)** Cohesion of the *URA3* locus in the absence of Eco1 and Wpl1. 3xmAID alleles were depleted during G1 arrest before release into M phase arrest when DNA circles were excised. Strains MSB203, MSB204, MSB205 and MSB206 were used.

When Eco1 was depleted the smallest obstacle, GFP-lacI alone, blocked cohesin translocation (Figure 6B). This unexpected result was due specifically to loss of Eco1 because it did not occur when DMSO was added instead of auxin (data not shown) or when Eco1 lacked the AID tag (Figure 6C). The blockade was also dependent on the synthetic barrier: when Eco1 was depleted from a strain lacking GFP-lacI accumulation of cohesin at the 5’lacO site was abolished (data not shown). It appears that Eco1, presumably through Smc3 acetylation, facilitates translocation past DNA-bound GFP-lacI.

To explore the phenomenon further, we examined cohesin translocation in a strain with an *smc1- D1164E* allele. This mutation diminishes ATP-hydrolysis by cohesin, creating chromatin-bound complexes that do not require Eco1 for protection against Wpl1 unloading (16, 51, 52). Importantly, this mutation reduces Smc3 acetylation by Eco1 (52). In this strain, *URA3* transcription during an M phase arrest also caused cohesin to accumulate at GFP-lacI alone (Figure 6D). These results reinforce the notion that acetylation of Smc3 by Eco1 influences how cohesin navigates the GFP-lacI array.

When Wpl1 was depleted, two notable changes occurred. First, depletion of the protein abolished the three mCherry barrier, yielding a slight increase in accumulation at the four mCherry barrier (Figure 6E). The small signal amplitude may be related to earlier observations that cohesin levels are lower at enrichment sites in the absence of Wpl1 (9, 17, 18). These data suggest that cohesin translocation is less restrained in the absence of Wpl1. The second notable change in figure 6E is that accumulation of cohesin at the four mCherry chimera occurred even in the absence of *URA3* induction. This feature is not unique to M phase-arrested cells. During late G1 arrest when cohesin accumulates at the four mCherry chimera, loss of Wpl1 also eliminated the requirement for *URA3* induction (Figure S6A). Misregulation of *URA3* in the absence of Wpl1 could explain the results. However, *URA3* transcription was found to be normal (Figure S5D and S6B). Instead, it seems that basal transcription of *URA3* is sufficient to cause cohesin accumulation at synthetic obstacles in the absence of Wpl1. If cohesin binds less specifically in Wpl1 mutants (53), and the residence time on DNA is longer (48), basal transcription of *URA3* might be sufficient to mobilize mis-targeted complexes toward downstream synthetic barriers.

When Eco1 and Wpl1 were co-depleted, cohesin bypassed the three mCherry chimera yet the complex was blocked efficiently by the larger four mCherry chimera (Figure 6F). Cohesin accumulation did not require *URA3* induction, as seen before when depleting Wpl1 alone. Interestingly, the paradoxical accumulation of cohesin at GFP-lacI in the absence of Eco1 was abolished when both proteins were co-depleted. Apparently, the impact of Wpl1 depletion is dominant when both proteins are absent. Perhaps with a longer DNA residence time conferred by Wpl1 loss, cohesin eventually bypasses barriers with fewer than four mCherries.

Interpretation of the cohesin translocation results in the absence of Eco1 and Wpl1 requires knowledge of the cohesive state of the complexes under study. Thus, we measured cohesion of the chromosomal domain in the absence of the regulators. For simplicity, the experiments were performed without transcriptional induction. Depletions were achieved during G1 arrest and then cells were grown to an M phase arrest before DNA circles were excised. As expected, depletion of Eco1 abolished cohesion (Figure 6G). Co-depletion of Eco1 and Wpl1 also yielded a severe cohesion defect, as anticipated from earlier work (See (54), and references there in). Wpl1 depletion was previously shown to have an intermediate effect on chromosomal cohesion (17, 18, 54). Here, depletion of Wpl1 abolished cohesion of the locus under study. These data show that the *URA3* gene is devoid of cohesive cohesin when Eco1 and or Wpl1 are depleted. Therefore, complexes that translocate under these conditions due so on only one chromatid even though both sisters are present.

## Discussion

In this study we show that synthetic barriers of sufficient size cause ectopic accumulation of the cohesin at synthetic roadblocks downstream of an inducible gene (Figure 2). That a DNA-bound obstacle built entirely of non-yeast proteins blocks the passage of cohesin shows that the complex translocates processively along DNA inside live cells. While our work was underway, a study using nuclease-dead Cas9 as a barrier also showed that cohesin moves processively *in vivo* (55). Our study goes further in three significant ways. First, we show that both cohesive and non-cohesive complexes translocate processively on chromosomes. Second, we show that the two types of complexes are blocked by barriers of different size (Figure 7A). Third, we show that stalled cohesive complexes block passage of non-cohesive complexes (Figure 7B).

**Figure 7.**
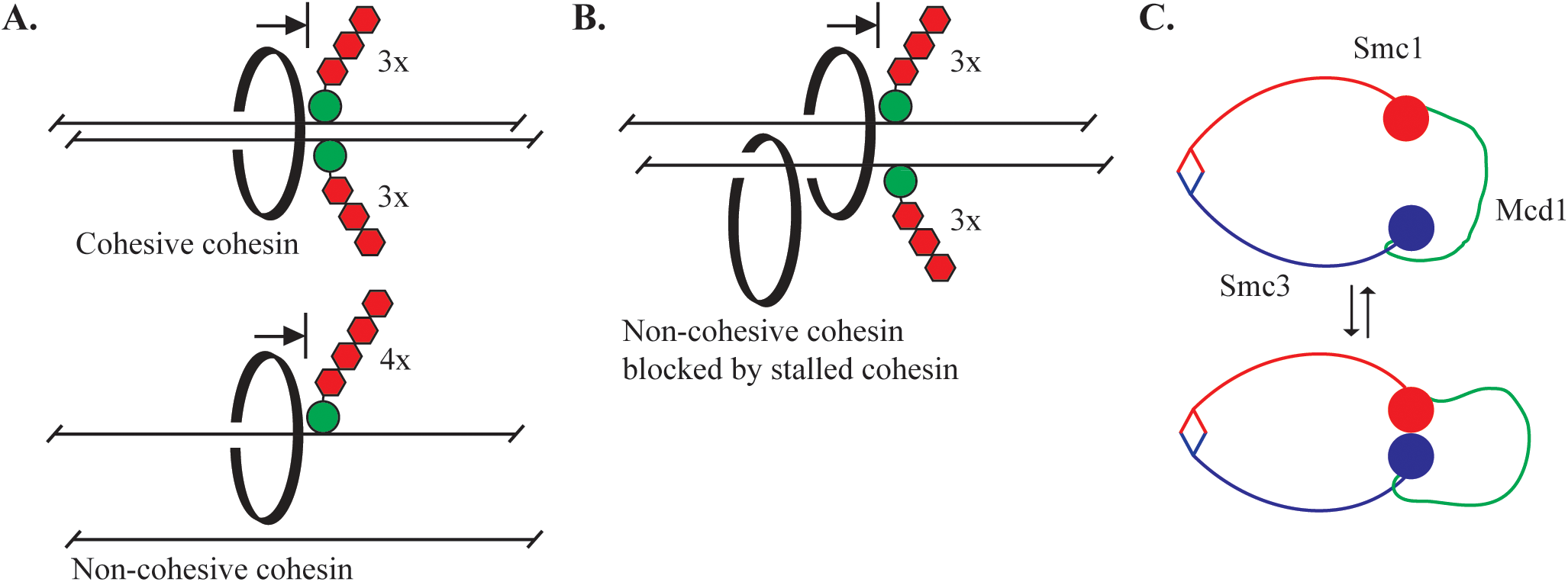
Processive translocation of cohesin *in vivo*. **A)** Cohesive cohesin cannot translocate past a barrier larger than three mCherries during M phase. Non-cohesive complexes translocating on single DNA (in late G1 or M phase) cannot translocate past a barrier larger than four mCherries. That cohesin is mobilized by transcription and then arrests at obstacles suggests that the complexes under study are bound topologically (see main text). **B)** Non-cohesive cohesin is blocked by cohesive cohesin that has stalled at a 3xmCherry barrier. **C)** The chambers of cohesin. The large topological compartment created by interactions of Smc3 and Smc1 with Mcd1 is bisected into two smaller chambers by engagement of the Smc3 and Smc1 heads (filled circles, (56)). The chamber(s) used during translocation have not yet been defined.

Cohesive cohesin has been shown to entrap both sister chromatids topologically within a single complex (3, 4). Presumably, the cohesive cohesin that translocates in our studies is similarly bound to DNA. Crosslinking studies have defined one large DNA chamber formed by the interactions of core subunits (Smc1, Smc3 and Mcd1), as well as two smaller DNA chambers partitioned from the larger chamber by the engaged heads of the Smc proteins (Figure 7C, (56)). These and additional cryo-EM studies have mapped cohesin within the chambers of cohesin (57, 58). Synthetic obstacles of increasing size serve as a molecular ruler of the embrace of cohesin through one or more of these chambers. Our work defines the upper size limit (measured in mCherries) for cohesin’s embrace of sister chromatids.

Non-cohesive cohesin can also bind single chromatids in a topological embrace but whether the non-cohesive complexes in our studies are similarly bound is less clear (3, 4, 59). Biochemical studies of cohesin translocation on single pieces of DNA have described active and passive movements. In the active case, cohesin moved linearly along DNA and extruded DNA loops in an ATP-dependent manner, the latter of which involved major conformational changes of the complex (27, 28, 60). This type of movement required the continual action of NIPBL/Scc2, a cofactor that stimulates the intrinsic cohesin ATPase activity (61). In congruence, loop extrusion *in vivo*, as well as less defined modes of transport, have been linked to NIPBL/Scc2 (33, 62, 63). For the following reasons, cohesin transport driven by NIPBL/Scc2 does not appear to involve topological entrapment of DNA. To begin with, loop extrusion *in vitro* was observed with crosslinked complexes that prevented DNA access to the central chamber (27). Moreover, a recent biochemical study showed that loop extruding complexes bypassed roadblocks that greatly exceeded the size of cohesin (64). Taken together, active movement of non-cohesive cohesin does not appear to involve topological engagement nor require transcription as a motive force.

In the passive movement case, topologically bound complexes were shown to travel in a diffusional manner on DNA *in vitro* (24, 25). Such complexes gained directional mobility when driven by transcribing RNA polymerase (24). Unlike loop extruding complexes, this type of movement was blocked by DNA bound obstacles. Given that the cohesin complexes in our studies were blocked by obstacles on DNA and that their movement required a transcriptional trigger, it is reasonable to assume that they are of the topologically bound variety. Whether NIPBL/Scc2 is recruited to complete cohesin transit all the way to a barrier, or whether non-coding transcription beyond the *URA3* gene contributes is still under investigation.

The threshold size of an impassible barrier changed depending on when cohesin was loaded onto chromosomes and when it translocated. Complexes loaded and transiting before S phase were blocked by a four mCherry obstacle (Figure 4). Non-cohesive complexes loaded after S phase were similarly blocked by a four mCherry barrier (Figure 5). Cohesive complexes, those which were loaded before S phase but translocated after, were blocked instead by a three mCherry barrier (figures 2 and 3). Collectively, these observations suggest that the cohesive state of a cohesin complex is the determinant of which mCherry barrier will block translocation. In this scenario, the requirement of larger barriers to block non-cohesive complexes may simply reflect the fact that these complexes entrap one less DNA duplex. In other words, non-cohesive complexes translocate past smaller barriers because their DNA chambers are filled with less DNA.

Both Eco1 and Wpl1 influence the size and distribution of chromosome loops in yeast, as do their homologs in higher eukaryotes (8, 9, 27, 32, 33, 65, 66). Whereas Wpl1 restricts loop size by limiting the residence time and location of cohesin on DNA, Eco1 is thought to restrict loop size through a mechanism(s) independent of cohesion establishment. A simple hypothesis holds that conditions which favor more extensive loop extrusion would also favor translocation past smaller synthetic barriers. Indeed, simultaneous depletion of Eco1 and Wpl1 or just Wpl1 alone increased the threshold size of a barrier in M phase cells from three mCherries to four mCherries (Figure 6E and 6F). However, we showed that the locus under study was not cohesed in the absence of Eco1 and Wpl1 (Figure 6G). Thus, a simpler explanation may prevail: the increase in barrier size from three to four mCherries reflects the absence of cohesive complexes in these mutants. This conclusion is entirely congruent with the cell cycle constraints described above.

Lastly, depletion of Eco1 alone yielded an entirely unexpected result: GFP-lacI alone was sufficient to impede cohesin passage (Figures 6B). The observation was reinforced with an *smc1- D1164E* mutant that hinders Smc3 acetylation by Eco1, as well as the intrinsic ATPase of cohesin (Figure 6D). Recent studies have shown that acetylation by Eco1 plays a role in anchoring the bases of cohesin-dependent loops (8, 65, 66). Whether the phenomenon is related to this paradoxical result will require further investigation.

We have shown that cohesive cohesin at smaller barriers blocks passage of topologically-bound non-cohesive complexes (Figure 5). A corollary to this conclusion is that sites of cohesin accumulation in genome mapping studies may consist of both cohesive non-cohesive cohesin. Recent mapping of cohesive cohesin in humans indicates that loop extrusion complexes do not bypass cohesive complexes but instead help position them (13). It has also been argued that stably bound cohesin complexes in yeast (possibly cohesive complexes) block passage of loop extruders (9). This raises the possibility that stalled cohesive cohesin in our assays also blocks passage of loop extruders. Maps of mammalian chromosomal loops often contain “stripes” that are thought to be a signature of translocating loop extrusion complexes (67). Fine structure mapping shows that stripes contain embedded dots, possibly from paused translocating complexes (68). The synthetic barriers described here may mimic the natural impediments (e.g. proteins bound at nucleosome depleted regions) that punctuate and terminate stripe patterns. Whether stalled cohesive cohesin complexes or topologically bound non-cohesive complexes are required for the punctuation of stripes remains to be determined.

## Materials and Methods

### Yeast strains and plasmids

Strains generated for this study were confirmed by PCR and/or DNA sequencing. Strains are listed in Table S2 and primers are listed in Table S1. GFP-lacI with a linked SV40 NLS was expressed from either a single or tandem integration of plasmid pGVH60, as noted in the strain table. These integrants were modified by PCR-mediated gene tagging using multimeric repeats of mCherry from plasmids pMAM12 and pMAM44 or three yeGFP(A206R)s from plasmid pMAM88. Strains with tandem pGVH60 thus yielded simultaneous expression a GFP-lacI chimera and GFP-lacI alone. Strains co-expressing GFP-lacI/GFP-lacI-mCherry chimeras, however, produced identical results to strains expressing the mCherry chimeras alone (compare figures 4C and S7). Integrating plasmid pXDA2-FFAT was assembled by *in vivo* recombination from pAFS144-FFAT and pXRA2 (69). pXDL2-GAL1p-Sic1(4m)-AID*-9xmyc was similarly assembled from pXRL2 (69), pRS306-GAL-Mt-4A-SIC1-HA-HIS6-Act (41) and pKan-AID*- 9myc (70). Integrating plasmid pXDnMX-GAL1p-Scc1(R180D,R268D)-TAP was assembled by in vivo recombination from YIplac128-GAL1p-Scc1(R180D,R268D)-3xHA (LEU2), pXDnMX (69), and a PCR fragment containing a TAP tag. Other genomic AID-tagged genes were constructed by PCR tagging with the 3xmAID-5xflag module from pST1933 (NBRP ID: BYP8880; (71)). OsTir1 was from integrated pTIR5 (*ADH1p-OsTIR1-CaADH1t,* Thomas Eng thesis, U.C. Berkeley) or integrated pMSB27 (*ADH1p-OsTIR1*(yeast codon optimized)), which was derived by *in vivo* recombination from pXRT1 (69) and pMK200 (NBRP ID: BYP7569).

### Cell growth and arrest

To arrest cells in M phase for ChIP-qPCR, methionine was added (Cf = 2mM) to mid-log cultures grown in SC-met to shut-off *MET3p-cdc20*. After 2.5 hours (roughly 80% of the cells adopted a dumbbell shape), cells were washed twice with water and resuspended in SC media lacking uracil to induce *URA3*. Two hours later the cultures were fixed for ChIP, according to (35). To arrest cells in M phase for analysis of cohesion, cultures were grown in SC-met plus dextrose for 6-8 hours and then diluted 200-400 fold into SC-met plus raffinose for overnight growth. When cells reached mid-log the following morning, methionine was added. 150 minutes later, cells were washed twice and resuspended in SC media lacking uracil. Two hours later, galactose was added (Cf = 2%) to induce circularization by the R recombinase. Cells were fixed with paraformaldehyde after two additional hours. To arrest cells for ChIP in late G1, dextrose grown cultures were diluted 200-400 fold into SC-met plus raffinose for overnight growth. When the cultures reached mid-log the following morning, alpha factor was added (Cf = 10 µM), and 90 minutes later galactose was added to induce expression of *GAL1p-SIC1(4m)-AID*-9xmyc*. After one additional hour, cultures were washed twice with water and resuspended in SC-met plus galactose and pronase E (Cf = 0.1 mg/ml) to release from pheromone arrest. One hour later, the cells were washed twice and resuspended in galactose lacking both methionine and uracil. Two hours later, the cultures were fixed for ChIP. For release from late G1 arrest, cells were grown into the arrest as described above but with the following exceptions. After cells were washed to remove alpha factor, they were resuspended in galactose media that lacked uracil but contained methionine and pronase E. Two hours later, dextrose (Cf = 2%) was added to stop expression of Sic1(4m)-AID*-9xmyc and 1-naphthaleneacetic acid (NAA, Cf = 100 µM) was added to degrade the existing protein. Two hours later when cells accumulated in M phase, cultures were fixed for ChIP. To express Mcd1-TAP exclusively in M phase, cells were arrested in M by depletion of Cdc20, as described above. Two and half hours later, galactose was added to induce expression of *GAL1p-MCD1(R180D,R268D)-TAP*. One hour later, NAA (or DMSO) was added to degrade the existing Mcd1-3xmAID. Thirty minutes later, the cells were washed twice with water and resuspended with the same media lacking uracil. Cells were fixed for ChIP after an additional two hours. For depletion of the cohesin regulators, strains with the various 3xmAID alleles were treated with alpha factor for 90 minutes before adding NAA for 60 additional minutes. After release from alpha factor arrest, cells were resuspended in media containing methionine for arrest in the subsequent M phase. 90 minutes later, cells were washed twice before resuspending with media lacking uracil. Two hours later, cells were fixed for ChIP. For cohesion assays involving protein depletions, cells were grown similarly except that cultures were diluted into SC-met plus raffinose for overnight growth. In addition, galactose was added once cells reached the M phase arrest and cells were fixed two hours later.

### Other techniques

ChIP-qPCR, RT-qPCR and fluorescence microscopy were performed as described in (35). For ChIP-qPCR, the mean and standard deviation of three or more biological replicates are presented, unless specified otherwise. All values were normalized to an unlinked cohesin-bound site (549.7 kb on Chr IV). Statistical significance was determined by pairwise Student’s t tests relative to the induced 0x mCherry strain, unless specified otherwise. Immunoblots were probed with anti-GFP (G1544, Sigma), anti-myc (9E10, Roche), anti-mini-AID (M214-3, MBL), anti- Pgk1 (ab113687, Abcam) or anti-Rad23 (from the Madura laboratory), and secondary antibodies (W401B and W402B, Promega). ChIP was performed with anti-TAP and protein A Dynabeads (CAB1001 and 10001D, Invitrogen). A ChemiDoc instrument (BioRad) was used for gel imaging.

## Acknowledgements

We thank the following individuals for reagents: Jason Brickner (pAFS144-FFAT), Michael Knop (pMAM12, pMAM44 and pMAM88), Kiran Madura (anti-Rad23), Adam Rudner (pTIR5), Benjamin Rowland (yeast BR423 *smc1-D1164E*) and Frank Uhlmann (YIplac128-GAL1p- Scc1(R180D,R268D)-3xHA (LEU2)). We thank Paul Kraycer for comments on the manuscript. This work was funded by United States Public Health Service Grant NIGMS 51402 (to M.R.G.) and a Busch Biomedical Grant from Rutgers University. The Flow Cytometry Core Facility of Robert Wood Johnson Medical School and The Rutgers Cancer Institute of New Jersey is supported by grants from the National Cancer Institute (Grant P30ES005022) and NIH (Grant S10OD026876).

---

**Figure S1.**
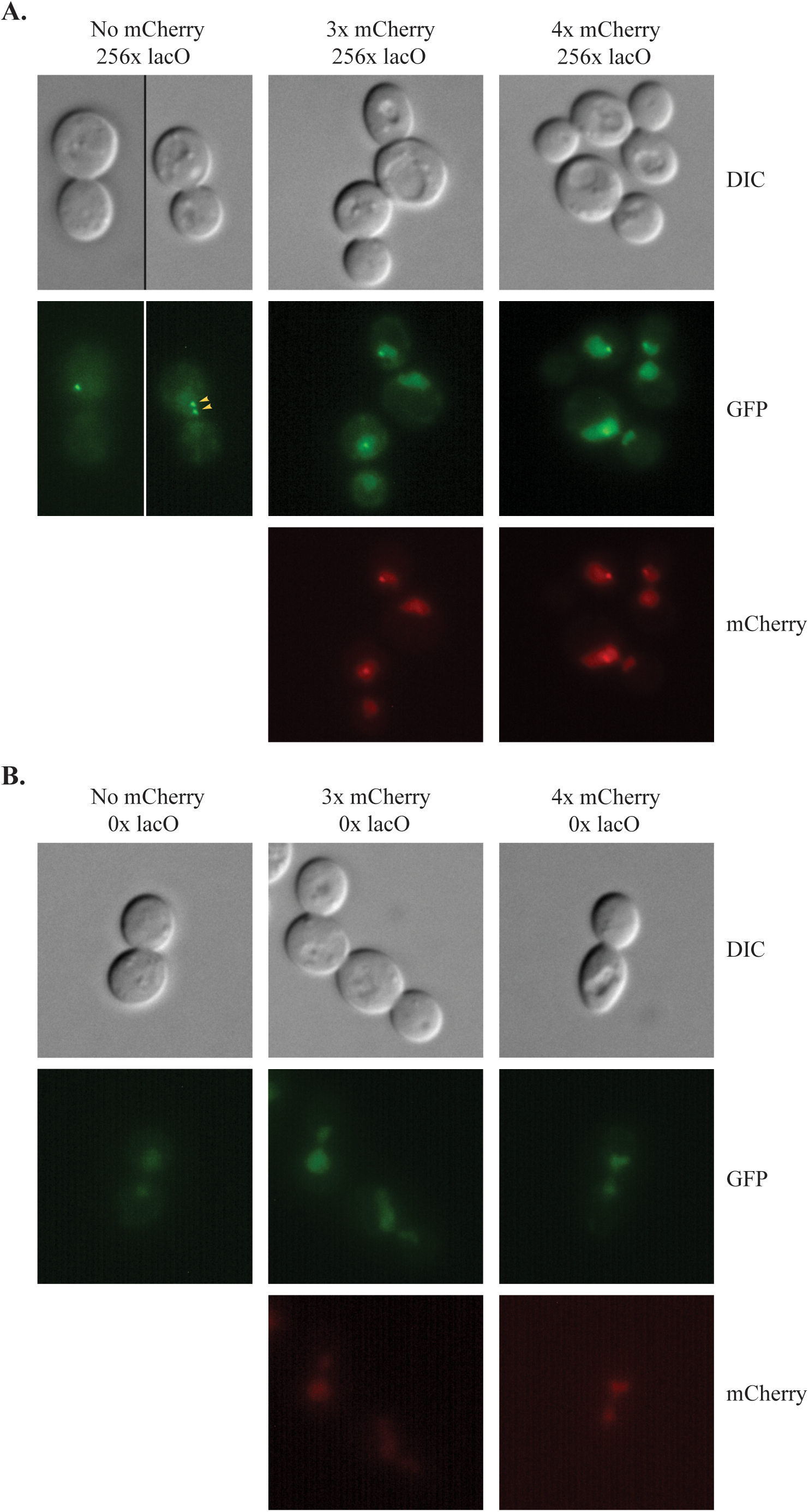
Representative images of cells expressing GFP-lacI chimeras. **A)** Strains MSB35.2, MSB167 and MRG7060 were photographed after M phase arrest. Fluorescent protein chimeras accumulated in the nucleus owing to a C-terminal SV40 NLS. Binding of GFP-lacI alone to the lacO array yielded green foci with little background fluorescence. When chromatids separated, two dots were seen (yellow arrows). Fusion of mCherries to GFP-lacI increased background nuclear fluorescence, suggesting that the C-terminal addition to lacI interferes with chimera binding. Nevertheless, sufficient binding occurred to yield discrete, albeit less intense foci. **B)** In the absence of a lacO array (strains MRG7372, MRG7373 and MRG7374), no foci were seen, indicating that GFP-lacI-mCherry chimeras do not intrinsically aggregate.

**Figure S2.**
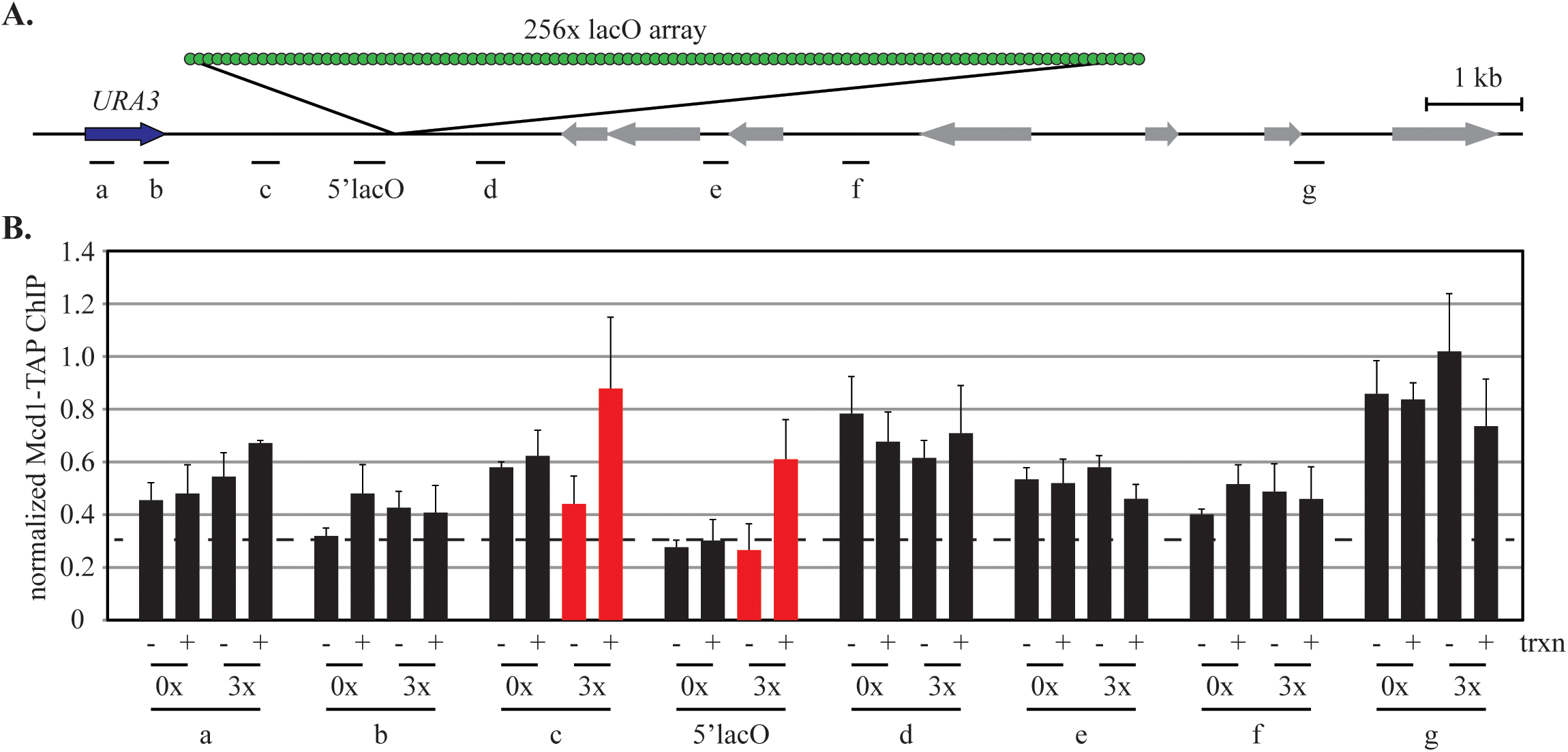
Cohesin distribution in strains expressing GFP-lacI-3xmCherry or GFP-lacI. **A)** A map of the chromosome IIIR telomeric region bearing the engineered locus. Sites that were analyzed are designated with lines and labels underneath the map. Drawn to scale. **B)** ChIP-qPCR of Mcd1-TAP. Strains MSB165.1 and MSB114 were used. Transcription was induced after M phase arrest.

**Figure S3.**
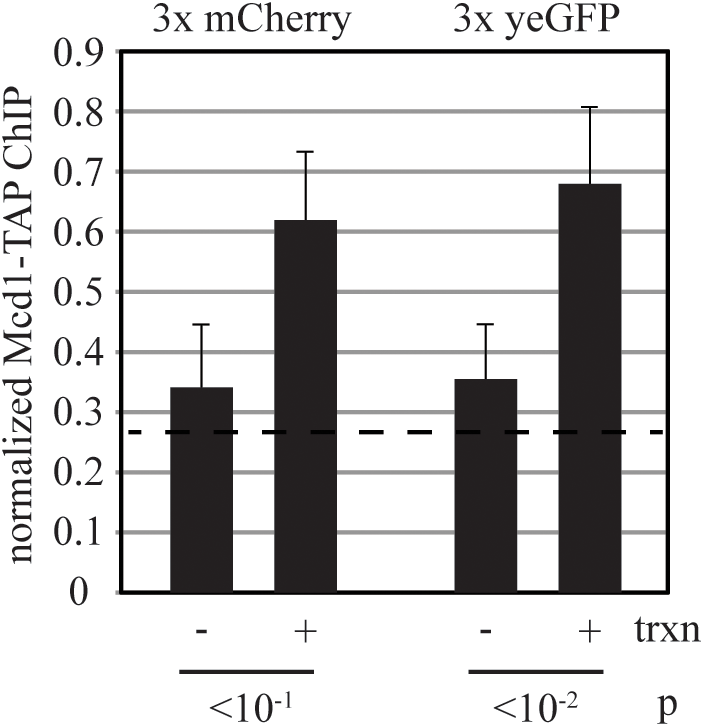
A barrier consisting of GFP-lacI plus three copies of yeGFP. Strains MSB165.1 and MRG7320 were used. Each of the three yeGFP multimers carries the A206R mutation that abolishes weak dimerization of GFP (73). Cultures were grown as described in figure 2. Three yeGFP(A206R) yielded the same result as three mCherries, indicating that residual multimerization of mCherry or some other mCherry artifact was not responsible for barrier activity. Two additional observations disfavor the notion that an aberrant property of mCherry is responsible for barrier activity: 1) different sized obstacles are required at different stages of the cell cycle (compare figures 2 and 4); 2) the timing of transcription determines the size of an effective obstacle (figures 2 and 3). Loci with notably high changes in cohesin accumulation are shown in red.

**Figure S4.**
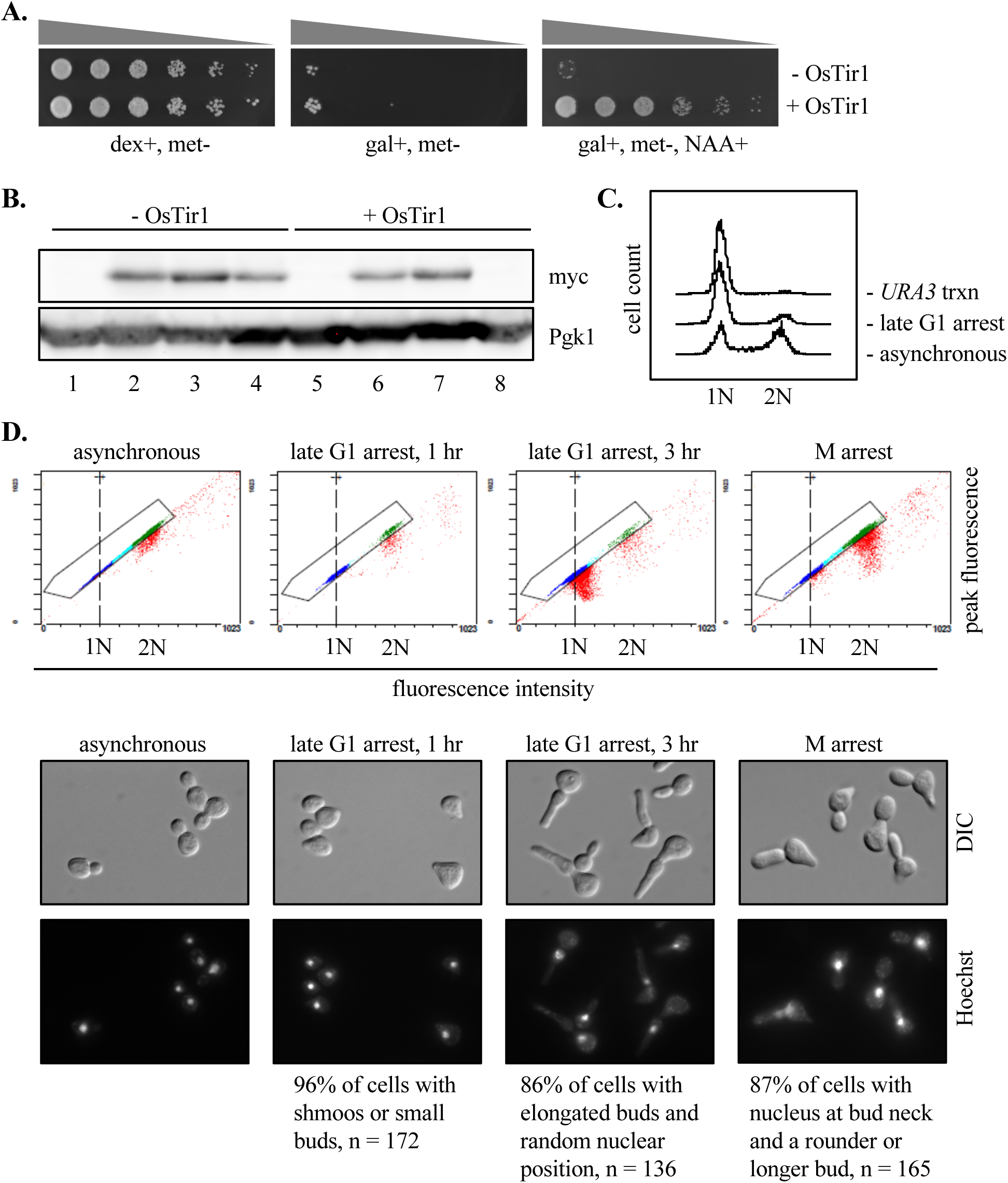
Stepwise cell cycle arrests using controlled expression of “non-degradable” Sic1. **A**) Cell viability following induction and clearance of Sic1(4m)-AID*-9xmyc. Strains bearing a *GAL1p-SIC1(4m)-AID*-9xmyc* allele (MRG7027 and MRG7033) were spotted in 10-fold serial dilutions on the indicated plates. Galactose-induced lethality due to Sic1(4m)-9xmyc-AID* expression was over-ridden by 100 µM 1-naphthaleneacetic acid (NAA), which triggers degradation of the AID* chimera by OsTir1 in strain MRG7033. Methionine was omitted from all plates for *MET3p-cdc20* expression. **B**) Validation of Sic1(4m) protein expression levels. The same strains were grown according to the arrest and release protocol described in the Materials and Methods. Lanes 1 and 5 = asynchronous growth in dextrose; lanes 2 and 6 = initial late G1 arrest in galactose media, one hour after removing alpha factor; lanes 3 and 7 = extended arrest at late G1 in galactose media, three hours after removing alpha factor; lanes 4 and 8 = two hours after adding NAA. Note that the myc-labelled Sic1(4m)-AID* band was cleared in lane 8 allowing this strain to advance to an M phase arrest. **C**) Validation of late G1 arrest by flow cytometry. Strain MRG7027 was grown into a permanent late G1 arrest as described in Materials and Methods, and used in figure 4B. DNA content was assessed in asynchronous culture, after late G1 arrest and then after subsequent *URA3* induction. **D**) Validation of sequential cell cycle arrests. Top row of panels - Dot plots of flow cytometry data (peak fluorescence vs fluorescence intensity). Strain MRG7033 was used. Cells with clear 1N and 2N DNA content are shown in navy blue and green, respectively. After a prolonged time in late G1 arrest, a large fraction of cells (red) fell outside the gated area due to an elongated bud morphology (due to Sic1-suppressed Cdk activity, (74)) that adversely affects peak fluorescence. Nevertheless, the data show that cells shown in red shifted from 1N to 2N DNA content during progression from the late G1 to M arrests. Bottom rows of panels - Direct visualization of cells and cell nuclei. The percentage of cells that met the specified morphological criteria of each cell cycle arrest are listed below the images. DNA was stained with Hoechst dye after ethanol fixation. n = number of cells examined.

**Figure S5.**
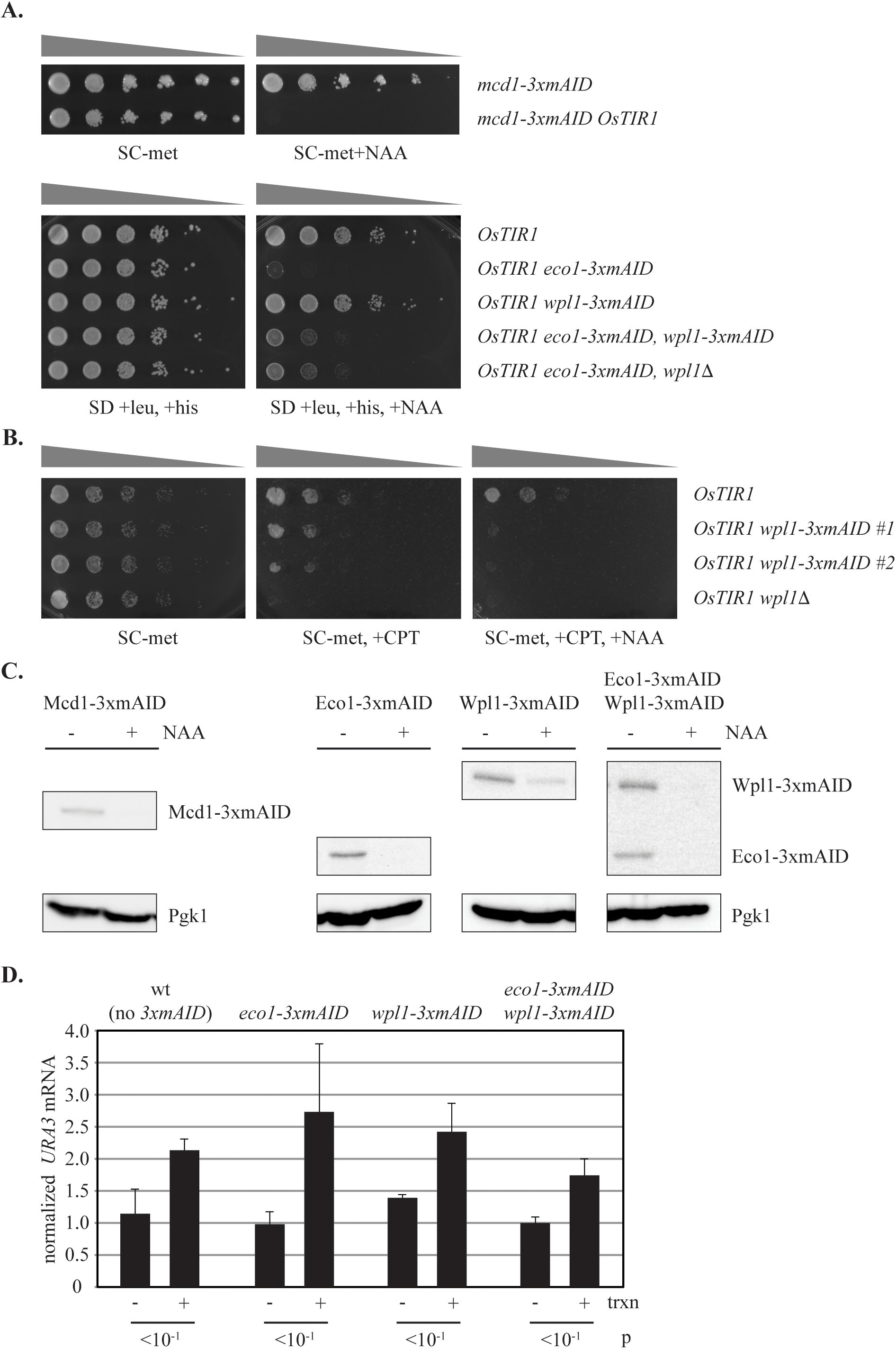
Depletion of Mcd1 and cohesin regulators. **A)** Cell viability following depletion of Mcd1, Eco1 and Wpl1. Strains MRG7312.1, MRG7317, MRG7270.1, MRG7301.1, MRG7327.1, MRG7332.1 and MRG7355.1 bearing the indicated *3xmAID* alleles and *OsTIR1* were spotted in 10-fold serial dilutions on selective plates containing or lacking 100 µM NAA. Depletion of Mcd1 or Eco1 caused loss of viability. Co-depletion of Eco1 and Wpl1 restored viability. Similar results were obtained whether Wpl1 was depleted or the *WPL1* gene was deleted, indicating that depletion phenocopied the null mutation. **B)** Camptothecin sensitivity following Wpl1 depletion. Strains MRG7269.1, MRG7329 (isolate 1), MRG7329 (isolate 2) and MRG6830.1 were spotted on SC-met containing or lacking 20 µg/ml camptothecin (CPT) and 100 µM NAA in 5-fold serial dilutions. Strains lacking Wpl1 were sensitive to the DNA damaging agent (See (53) and references therein). The middle panel shows that tagging with 3xmAID slightly diminished the function of Wpl1. The right panel shows that depletion of the protein with NAA phenocopied the null mutation. **C)** Depletion of 3xmAID-tagged proteins. Strains MRG7317, MRG7301.1, MRG7327.1, and MRG7332.1 were used. Cells were grown to mid-log in SC-met and treated with either 100 µM NAA or DMSO for 10 minutes for Mcd1 depletion and 60 minutes for the others. The tagged proteins were immunoblotted with an antibody to mAID. **D)** RT-qPCR analysis of *URA3* mRNA following depletion of Eco1 and Wpl1. Strains MRG7270.1, MRG7301.1, MRG7327.1, and MRG7332.1 were grown according to the protocol described in figure 5A. Transcript levels were measured relative to *ACT1* mRNA and normalized to a single trial of the uninduced wt control. The data show that loss of Wpl1 and or Eco1 did not perturb normal *URA3* induction.

**Figure S6.**
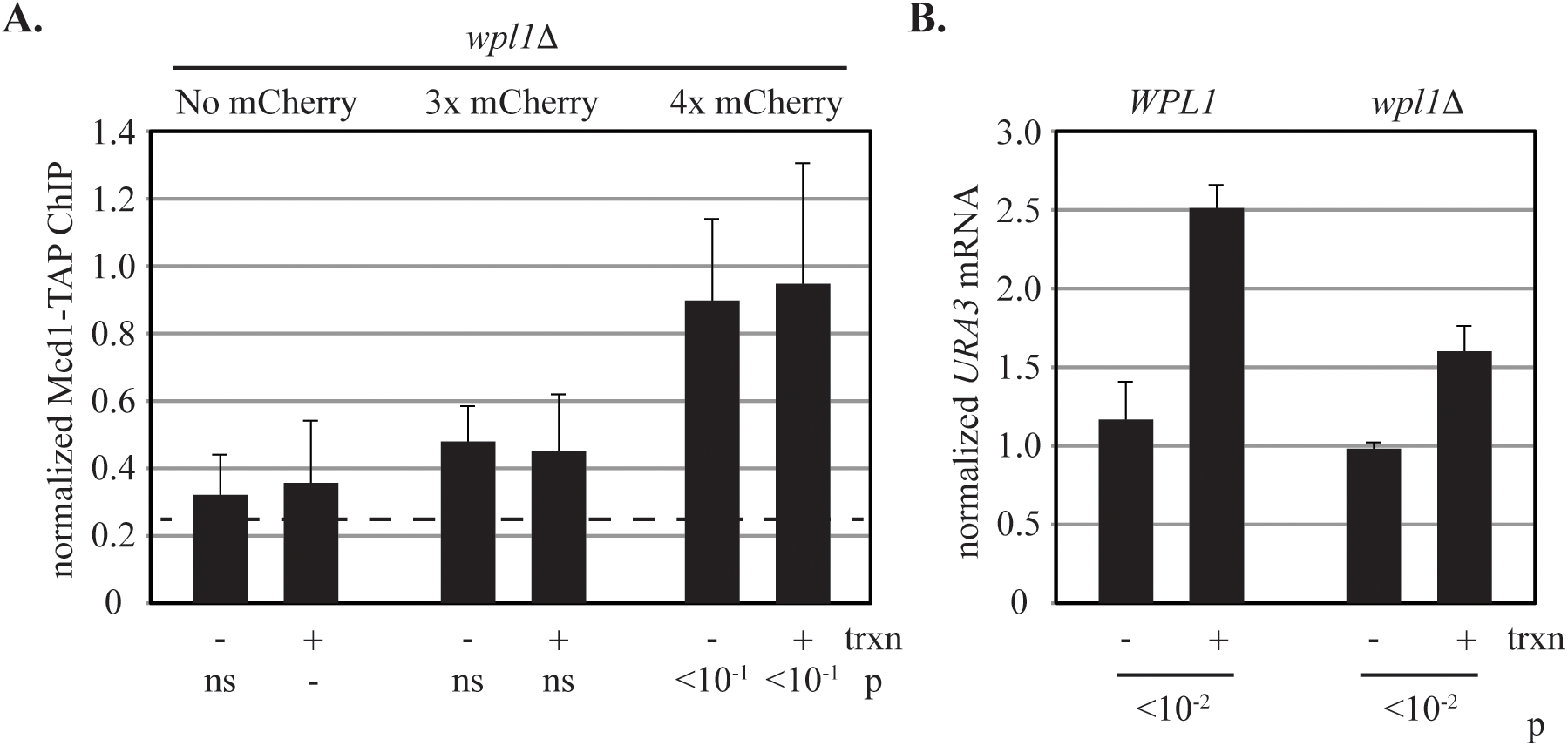
A synthetic barrier to cohesin translocation in late G1-arrested *wpl1Δ* cells. **A)** ChIP-qPCR of Mcd1-TAP. Strains MRG7235.1, MRG7232.1 and MRG7238.1 were grown according to the protocol described in figure 4A. Loss of Wpl1 caused accumulation of cohesin at the four mCherry barrier without *URA3* transcriptional induction. **B)** RT-qPCR analysis of *URA3* mRNA levels. Strains MRG7027 and MRG7235.1 were grown asynchronously before RNA extracts were prepared. Values normalized as in Figure S4. Loss of Wpl1 did not alter the basal level of *URA3* transcription or the ability to induce the gene.

**Figure S7.**
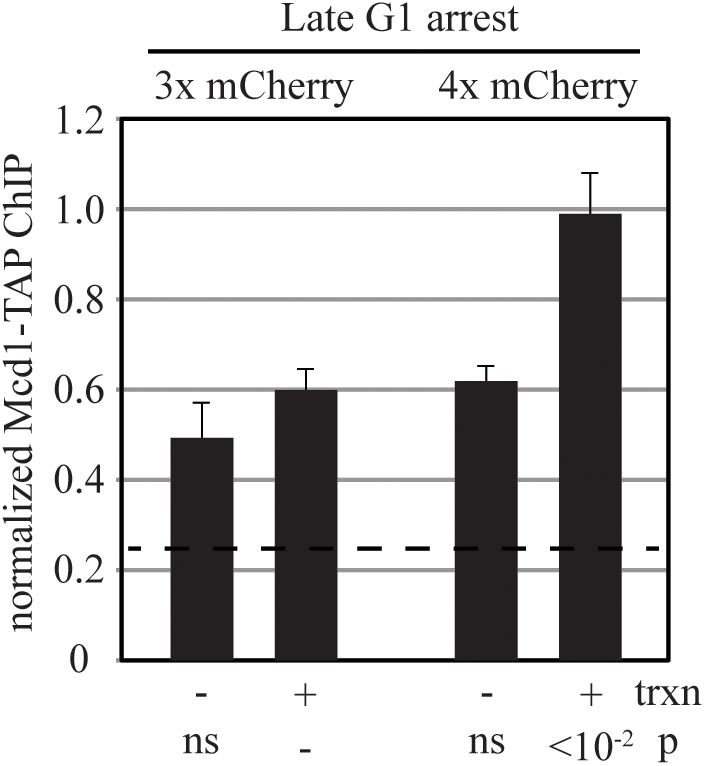
Barrier activity in strains expressing homogeneous GFP-lacI-mCherry chimeras. *URA3* was induced in strains MRG7115 and MRG7122 after arrest in late G1 by Sic1(4m)- AID*-9xmyc. Binding of cohesin at 5’lacO was analyzed by ChIP-qPCR of Mcd1-TAP. Barrier function required four mCherries, similar to the results found with strains co-expressing the mCherry chimeras along with GFP-lacI alone (Figure 4).

**Table S1.** Yeast strains.

| Name | Genotype | Parent | Ref | Fig. |
| --- | --- | --- | --- | --- |
| MSB35.2 | *MATa hmr-git1Δ::RS-URA3f-loxP-256xlacO-RS ade2-1::(HIS3p-GFP-lacI::ADE2), leu2-3,112::(GAL1p-RecR::LEU2), cdc20::MET3p-cdc20::loxP can1-100 his3-11,15 trp1-1 ura3-1* | MSB31 | Borrie et al 2017 | 3C, S1A |
| MSB114 | *MATa hmr-git1Δ::RS-URA3f-loxP-256xlacO-RS ade2-1::(HIS3p-GFP-lacI::ADE2), leu2-3,112::(GAL1p-RecR::LEU2), cdc20::MET3p-cdc20::loxP MCD1-TAP::natMX ura3Δ::hphMX can1-100 his3-11,15 trp1-1* | MSB35.2 | This study | 1C, 2E, S2B |
| MRG6947 | *MATa hmr-git1Δ::RS-URA3f-loxP-256xlacO-RS leu2-3,112::(GAL1p-RecR::LEU2), MCD1-TAP::natMX cdc20::MET3p-cdc20::loxP his3-11,15::HIS3p-GFP-FFAT-lacI::ADE2::HIS3p-GFP-FFAT-lacI::HIS3 ura3Δ::AgTEFp-SpHIS5 ade2-1 can1-100 trp1-1* | MRG6767 x MRG6900 | “” | 1C |
| MSB174 | *MATa hmr-git1Δ::RS-URA3f-loxP-256xlacO-RS ade2-1::HIS3p-GFP-lacI::ADE2 leu2-3,112::(GAL1p-RecR::LEU2), ura3Δ::AgTEFp-SpHIS5 bar1Δ can1-100 his3-11,15 trp1-1* | MRG6860 | “” | 2C |
| MSB179 | *MATa hmr-git1Δ::RS-URA3f-loxP-256xlacO-RS ade2-1::HIS3p-GFP-lacI-1xmCherry::hphMX::ADE2 leu2-3,112::(GAL1p-RecR::LEU2), ura3Δ::AgTEFp-SpHIS5 bar1Δ can1-100 his3-11,15 trp1-1* | MSB174 | “” | 2C |
| MSB180 | *MATa hmr-git1Δ::RS-URA3f-loxP-256xlacO-RS ade2-1::HIS3p-GFP-lacI-2xmCherry::hphMX::ADE2 leu2-3,112::(GAL1p-RecR::LEU2), ura3Δ::AgTEFp-SpHIS5 bar1Δ can1-100 his3-11,15 trp1-1* | MSB174 | “” | 2C |
| MSB181 | *MATa hmr-git1Δ::RS-URA3f-loxP-256xlacO-RS ade2-1::HIS3p-GFP-lacI-3xmCherry::hphMX::ADE2 leu2-3,112::(GAL1p-RecR::LEU2), ura3Δ::AgTEFp-SpHIS5 bar1Δ can1-100 his3-11,15 trp1-1* | MSB174 | “” | 2C |
| MRG7102 | *MATa hmr-git1Δ::RS-URA3f-loxP-256xlacO-RS ade2-1::HIS3p-GFP-lacI-4xmCherry::kanMX::ADE2 leu2-3,112::(GAL1p-RecR::LEU2), ura3Δ::AgTEFp-SpHIS5 bar1Δ can1-100 his3-11,15 trp1-1* | MSB181 | “” | 2C |
| MRG6810 | *MATa hmr-git1Δ::RS-URA3f-loxP-256xlacO-RS ade2-1::HIS3p-GFP-lacI-* | MSB164 | “” | 2E |
|  | *1xmCherry::hphMX::ADE2::HIS3p-GFP-lacI::ADE2 leu2-3,112::(GAL1p-RecR::LEU2), cdc20::MET3p-cdc20::loxP MCD1-TAP::natMX ura3Δ::AgTEFp-SpHIS5 can1-100 his3-11,15 trp1-1* |  |  |  |
| MRG6812 | *MATa hmr-git1Δ::RS-URA3f-loxP-256xlacO-RS ade2-1::HIS3p-GFP-lacI-2xmCherry::hphMX::ADE2::HIS3p-GFP-lacI::ADE2 leu2-3,112::(GAL1p-RecR::LEU2), loxP::MET3p-cdc20 MCD1-TAP::natMX ura3Δ::AgTEFp-SpHIS5 can1-100 his3-11,15 trp1-1* | MSB164 | “” | 2E |
| MSB165.1 | *MATa hmr-git1Δ::RS-URA3f-loxP-256xlacO-RS ade2-1::HIS3p-GFP-lacI-3xmCherry::hphMX::ADE2::HIS3p-GFP-lacI::ADE2 leu2-3,112::(GAL1p-RecR::LEU2), cdc20::MET3p-cdc20::loxP MCD1-TAP::natMX ura3Δ::AgTEFp-SpHIS5 can1-100 his3-11,15 trp1-1* | MSB164 | “” | 2E,<br>S2B,<br>S3 |
| MRG7060 | *MATa hmr-git1Δ::RS-URA3f-loxP-256xlacO-RS ade2-1::HIS3p-GFP-lacI-4xmCherry::kanMX::ADE2::HIS3p-GFP-lacI::ADE2 leu2-3,112::(GAL1p-RecR::LEU2), cdc20::MET3p-cdc20::loxP MCD1-TAP::natMX ura3Δ::AgTEFp-SpHIS5 can1-100 his3-11,15 trp1-1* | MSB165.1 | “” | 2E,<br>3C,<br>S1A |
| MRG6806 | *MATa hmr-git1Δ::RS-URA3f-loxP-256xlacO-RS ade2-1::HIS3p-GFP-lacI-1xmCherry::hphMX::ADE2::HIS3p-GFP-lacI::ADE2 leu2-3,112::(GAL1p-RecR::LEU2), cdc20::MET3p-cdc20::loxP can1-100 his3-11,15 trp1-1 ura3-1* | MSB35.2 | “” | 3C |
| MRG6808 | *MATa hmr-git1Δ::RS-URA3f-loxP-256xlacO-RS ade2-1::HIS3p-GFP-lacI-2xmCherry::hphMX::ADE2::HIS3p-GFP-lacI::ADE2 leu2-3,112::(GAL1p-RecR::LEU2), cdc20::MET3p-cdc20::loxP can1-100 his3-11,15 trp1-1 ura3-1* | MSB35.2 | “” | 3C |
| MSB167 | *MATa hmr-git1Δ::RS-URA3f-loxP-256xlacO-RS ade2-1::HIS3p-GFP-lacI-3xmCherry::hphMX::ADE2::HIS3p-GFP-lacI::ADE2 leu2-3,112::(GAL1p-RecR::LEU2), cdc20::MET3p-cdc20::loxP can1-100 his3-11,15 trp1-1 ura3-1* | MSB165.1<br>x<br>W303-1B | “” | 3C,<br>S1A |
| MRG7027 | *MATa hmr-git1Δ::RS-URA3f-loxP-256xlacO-RS ade2-1::(HIS3p-GFP-lacI::ADE2), leu2-3,112::GAL1p-SIC1(4m)-AID\*-9xmyc::LEU2 cdc20::MET3p-cdc20::loxP MCD1-TAP::natMX* | MRG7013 | “” | 4B,<br>4C,<br>S4A,<br>S4B, |
|  | *ura3Δ::AgTEFp-SpHIS5 bar1Δ can1-100 his3-11,15 trp1-1* |  |  | S4C,<br>S7B |
| MRG7026 | *MATa hmr-git1Δ::RS-URA3f-loxP-256xlacO-RS ade2-1::HIS3p-GFP-lacI-3xmCherry::hphMX::ADE2::HIS3p-GFP-lacI::ADE2 leu2-3,112::GAL1p-SIC1(4m)-AID\*-9xmyc::LEU2 cdc20::MET3p-cdc20::loxP MCD1-TAP::natMX ura3Δ::AgTEFp-SpHIS5 bar1Δ can1-100 his3-11,15 trp1-1* | MRG7015 | “” | 4B,<br>4C |
| MRG7059 | *MATa hmr-git1Δ::RS-URA3f-loxP-256xlacO-RS ade2-1::HIS3p-GFP-lacI-4xmCherry::kanMX::ADE2::HIS3p-GFP-lacI::ADE2 leu2-3,112::GAL1p-SIC1(4m)-AID\*-9xmyc::LEU2 cdc20::MET3p-cdc20::loxP MCD1-TAP::natMX ura3Δ::AgTEFp-SpHIS5 bar1Δ can1-100 his3-11,15 trp1-1* | MRG7026 | “” | 4B,<br>4C |
| MRG7033 | *MATa hmr-git1Δ::RS-URA3f-loxP-256xlacO-RS ade2-1::(HIS3p-GFP-lacI::ADE2), leu2-3,112::GAL1p-SIC1(4m)-AID\*-9xmyc::LEU2 cdc20::MET3p-cdc20::loxP MCD1-TAP::natMX trp1-1::CgTRP1-ADH1p-OsTIR1-CaADH1t ura3Δ::AgTEFp-SpHIS5 bar1Δ can1-100 his3-11,15* | MRG7027 | “” | 4D,<br>S4A,<br>S4B,<br>S4D |
| MRG7032 | *MATa hmr-git1Δ::RS-URA3f-loxP-256xlacO-RS ade2-1::HIS3p-GFP-lacI-3xmCherry::hphMX::ADE2::HIS3p-GFP-lacI::ADE2 leu2-3,112::GAL1p-SIC1(4m)-AID\*-9xmyc::LEU2 cdc20::MET3p-cdc20::loxP MCD1-TAP::natMX trp1-1::CgTRP1-ADH1p-OsTIR1-CaADH1t ura3Δ::AgTEFp-SpHIS5 bar1Δ can1-100 his3-11,15* | MRG7026 | “” | 4D |
| MRG7282 | *MATa hmr-git1Δ::RS-URA3f-loxP-256xlacO-RS ade2-1::HIS3p-GFP-lacI-4xmCherry::kanMX::ADE2::HIS3p-GFP-lacI::ADE2 leu2-3,112::GAL1p-SIC1(4m)-AID\*-9xmyc::LEU2 cdc20::MET3p-cdc20::loxP MCD1-TAP::natMX trp1-1::CgTRP1-ADH1p-OsTIR1-CaADH1t ura3Δ::AgTEFp-SpHIS5 bar1Δ can1-100 his3-11,15* | MRG7059 | “” | 4D |
| MRG6989 | *MATa hmr-git1Δ::RS-URA3f-loxP-256xlacO-RS ade2-1::(HIS3p-GFP-lacI::ADE2), cdc20::MET3p-cdc20::loxP ho::GAL1p-MCD1(R180D,R268D)-TAP::natMX::ho ura3Δ::AgTEFp-SpHIS5 bar1Δ can1-100 his3-11,15 leu2-3,112 trp1-1* | MRG6974<br>x<br>MRG6963 | “” | 5B |
| MRG7294 | *MATa hmr-git1Δ::RS-URA3f-loxP-256xlacO-RS ade2-1::HIS3p-GFP-lacI-* | MRG7290 | “” | 5B |
|  | *3xmCherry::hphMX::ADE2::HIS3p-GFP-lacI::ADE2 cdc20::MET3p-cdc20::loxP ho::GAL1p-MCD1(R180D,R268D)-TAP::natMX::ho ura3Δ::AgTEFp-SpHIS5 bar1Δ can1-100 his3-11,15 leu2-3,112 trp1-1* |  |  |  |
| MRG7292 | *MATa hmr-git1Δ::RS-URA3f-loxP-256xlacO-RS ade2-1::HIS3p-GFP-lacI-4xmCherry::kanMX::ADE2::HIS3p-GFP-lacI::ADE2 cdc20::MET3p-cdc20::loxP ho::GAL1p-MCD1(R180D,R268D)-TAP::natMX::ho ura3Δ::AgTEFp-SpHIS5 bar1Δ can1-100 his3-11,15 leu2-3,112 trp1-1* | MRG7288 | “” | 5B |
| MRG7317.1 | *MATa hmr-git1Δ::RS-URA3f-loxP-256xlacO-RS ade2-1::(HIS3p-GFP-lacI::ADE2), ho::GAL1p-MCD1(R180D,R268D)-TAP::natMX::ho mcd1-3xmAID-5xflag::bleMX trp1-1::CgTRP1-ADH1p-OsTIR1-CaADH1t cdc20::MET3p-cdc20::loxP ura3Δ::AgTEFp-SpHIS5 bar1Δ can1-100 his3-11,15 leu2-3,112* | MRG7312.1 | “” | 5C, S5A S5C |
| MRG7319.1 | *MATa hmr-git1Δ::RS-URA3f-loxP-256xlacO-RS ade2-1::HIS3p-GFP-lacI-3xmCherry::hphMX::ADE2::HIS3p-GFP-lacI::ADE2 ho::GAL1p-MCD1(R180D,R268D)-TAP::natMX::ho mcd1-3xmAID-5xflag::bleMX trp1-1::CgTRP1-ADH1p-OsTIR1-CaADH1t cdc20::MET3p-cdc20::loxP ura3Δ::AgTEFp-SpHIS5 bar1Δ can1-100 his3-11,15 leu2-3,112* | MRG7316.1 | “” | 5C |
| MRG7318.1 | *MATa hmr-git1Δ::RS-URA3f-loxP-256xlacO-RS ade2-1::HIS3p-GFP-lacI-4xmCherry::kanMX::ADE2::HIS3p-GFP-lacI::ADE2 ho::GAL1p-MCD1(R180D,R268D)-TAP::natMX::ho mcd1-3xmAID-5xflag::bleMX trp1-1::CgTRP1-ADH1p-OsTIR1-CaADH1t cdc20::MET3p-cdc20::loxP ura3Δ::AgTEFp-SpHIS5 bar1Δ can1-100 his3-11,15 leu2-3,112* | MRG7315.1 | “” | 5C |
| MRG7301.1 | *MATa hmr-git1Δ::RS-URA3f-loxP-256xlacO-RS ade2-1::(HIS3p-GFP-lacI::ADE2), cdc20::MET3p-cdc20::loxP MCD1-TAP::natMX trp1-1::(ADH1p-OsTIR1(codon optimized)::TRP1), eco1-3xmAID-5xflag::bleMX ura3Δ::AgTEFp-SpHIS5 bar1Δ can1-100 his3-11,15 leu2-3,112* | MRG7270.1 | “” | 6B, S5A, S5C, S5D |
| MRG7300.1 | *MATa hmr-git1Δ::RS-URA3f-loxP-256xlacO-RS ade2-1::HIS3p-GFP-lacI-3xmCherry::hphMX::ADE2::HIS3p-GFP-lacI::ADE2 cdc20::MET3p-cdc20::loxP MCD1-* | MRG7269.1 | “” | 6B |
|  | *TAP::natMX trp1-1::(ADH1p-OsTIR1(codon optimized)::TRP1), eco1-3xmAID-5xflag::bleMX ura3Δ::AgTEFp-SpHIS5 bar1Δ can1-100 his3-11,15 leu2-3,112* |  |  |  |
| MRG7302.1 | *MATa hmr-git1Δ::RS-URA3f-loxP-256xlacO-RS ade2-1::HIS3p-GFP-lacI-4xmCherry::kanMX::ADE2::HIS3p-GFP-lacI::ADE2 cdc20::MET3p-cdc20::loxP MCD1-TAP::natMX trp1-1::(ADH1p-OsTIR1(codon optimized)::TRP1), eco1-3xmAID-5xflag::bleMX ura3Δ::AgTEFp-SpHIS5 bar1Δ can1-100 his3-11,15 leu2-3,112* | MRG7275.1 | “” | 6B |
| MRG7270.1 | *MATa hmr-git1Δ::RS-URA3f-loxP-256xlacO-RS ade2-1::(HIS3p-GFP-lacI::ADE2), cdc20::MET3p-cdc20::loxP MCD1-TAP::natMX trp1-1::(ADH1p-OsTIR1(codon optimized)::TRP1), ura3Δ::AgTEFp-SpHIS5 bar1Δ can1-100 his3-11,15 leu2-3,112* | MRG6964<br>X<br>MRG7258.2 | “” | 6C,<br>S5A,<br>S5D |
| MRG7269.1 | *MATa hmr-git1Δ::RS-URA3f-loxP-256xlacO-RS ade2-1::HIS3p-GFP-lacI-3xmCherry::hphMX::ADE2::HIS3p-GFP-lacI::ADE2 cdc20::MET3p-cdc20::loxP MCD1-TAP::natMX trp1-1::(ADH1p-OsTIR1(codon optimized)::TRP1), ura3Δ::AgTEFp-SpHIS5 bar1Δ can1-100 his3-11,15 leu2-3,112* | MRG6964<br>X<br>MRG7258.2 | “” | 6C,<br>S5B |
| MRG7275.1 | *MATa hmr-git1Δ::RS-URA3f-loxP-256xlacO-RS ade2-1::HIS3p-GFP-lacI-4xmCherry::kanMX::ADE2::HIS3p-GFP-lacI::ADE2 cdc20::MET3p-cdc20::loxP MCD1-TAP::natMX trp1-1::(ADH1p-OsTIR1(codon optimized)::TRP1), ura3Δ::AgTEFp-SpHIS5 bar1Δ can1-100 his3-11,15 leu2-3,112* | MRG7240.1<br>X<br>MRG7258.2 | “” | 6C |
| MRG7151 | *MATa hmr-git1Δ::RS-URA3f-lacOPs-RS ade2-1::(HIS3p-GFP-lacI::ADE2), cdc20::MET3p-cdc20::loxP leu2-3,112::GAL1p-SIC1(4m)-AID\*-9xmyc::LEU2 MCD1-TAP::natMX ura3Δ::AgTEFp-SpHIS5 smc1-D1164E bar1Δ can1-100 his3-11,15 trp1-1* | MRG6792<br>X<br>MRG7027 | “” | 6D |
| MRG7146 | *MATa hmr-git1Δ::RS-URA3f-lacOPs-RS ade2-1::HIS3p-GFP-lacI-3xmCherry::hphMX::ADE2::HIS3p-GFP-lacI::ADE2 cdc20::MET3p-cdc20::loxP leu2-3,112::GAL1p-SIC1(4m)-AID\*-9xmyc::LEU2 MCD1-TAP::natMX ura3Δ::AgTEFp-SpHIS5 smc1-D1164E bar1Δ can1-100 his3-11,15 trp1-1* | MRG6887<br>X<br>MRG7130 | “” | 6D |
| MRG7159 | *MATa hmr-git1Δ::RS-URA3f-lacOPs-RS ade2-1::HIS3p-GFP-lacI-4xmCherry::kanMX::ADE2::HIS3p-GFP-lacI::ADE2 cdc20::MET3p-cdc20::loxP leu2-3,112::GAL1p-SIC1(4m)-AID\*-9xmyc::LEU2 MCD1-TAP::natMX ura3Δ::AgTEFp-SpHIS5 smc1-D1164E bar1Δ can1-100 his3-11,15 trp1-1* | MRG7146<br>X<br>MRG7150 | “” | 6D |
| MRG7327.1 | *MATa hmr-git1Δ::RS-URA3f-loxP-256xlacO-RS ade2-1::(HIS3p-GFP-lacI::ADE2), cdc20::MET3p-cdc20::loxP MCD1-TAP::natMX trp1-1::(ADH1p-OsTIR1(codon optimized)::TRP1), wpl1-3xmAID-5xflag::bleMX ura3Δ::AgTEFp-SpHIS5 bar1Δ can1-100 his3-11,15 leu2-3,112* | MRG7270.1 | “” | 6E,<br>S5A,<br>S5C,<br>S5D |
| MRG7329.1 | *MATa hmr-git1Δ::RS-URA3f-loxP-256xlacO-RS ade2-1::HIS3p-GFP-lacI-3xmCherry::hphMX::ADE2::HIS3p-GFP-lacI::ADE2 cdc20::MET3p-cdc20::loxP MCD1-TAP::natMX trp1-1::(ADH1p-OsTIR1(codon optimized)::TRP1), wpl1-3xmAID-5xflag::bleMX ura3Δ::AgTEFp-SpHIS5 bar1Δ can1-100 his3-11,15 leu2-3,112* | MRG7269.1 | “” | 6E,<br>S5B |
| MRG7324.1 | *MATa hmr-git1Δ::RS-URA3f-loxP-256xlacO-RS ade2-1::HIS3p-GFP-lacI-4xmCherry::kanMX::ADE2::HIS3p-GFP-lacI::ADE2 cdc20::MET3p-cdc20::loxP MCD1-TAP::natMX trp1-1::(ADH1p-OsTIR1(codon optimized)::TRP1), wpl1-3xmAID-5xflag::bleMX ura3Δ::AgTEFp-SpHIS5 bar1Δ can1-100 his3-11,15 leu2-3,112* | MRG7275.1 | “” | 6E |
| MRG7332.1 | *MATa hmr-git1Δ::RS-URA3f-loxP-256xlacO-RS ade2-1::(HIS3p-GFP-lacI::ADE2), cdc20::MET3p-cdc20::loxP MCD1-TAP::natMX trp1-1::(ADH1p-OsTIR1(codon optimized)::TRP1), wpl1-3xmAID-5xflag::bleMX eco1-3xmAID-5xflag::kanMX ura3Δ::AgTEFp-SpHIS5 bar1Δ can1-100 his3-11,15 leu2-3,112* | MRG7327.1 | “” | 6F,<br>S5A,<br>S5C,<br>S5D |
| MRG7356.1 | *MATa hmr-git1Δ::RS-URA3f-loxP-256xlacO-RS ade2-1::HIS3p-GFP-lacI-3xmCherry::hphMX::ADE2::HIS3p-GFP-lacI::ADE2 cdc20::MET3p-cdc20::loxP MCD1-TAP::natMX trp1-1::(ADH1p-OsTIR1(codon optimized)::TRP1), wpl1-3xmAID-5xflag::bleMX eco1-3xmAID-5xflag::kanMX ura3Δ::AgTEFp-SpHIS5 bar1Δ can1-100 his3-11,15 leu2-3,112* | MRG7329.1 | “” | 6F |
| MRG7353.1 | *MATa hmr-git1Δ::RS-URA3f-loxP-256xlacO-RS* *ade2-1::HIS3p-GFP-lacI-4xmCherry::hphMX::ADE2::HIS3p-GFP-lacI::ADE2 cdc20::MET3p-cdc20::loxP MCD1-TAP::natMX trp1-1::(ADH1p-OsTIR1(codon optimized)::TRP1), wpl1-3xmAID-5xflag::bleMX eco1-3xmAID-5xflag::kanMX ura3Δ::AgTEFp-SpHIS5 bar1Δ can1-100 his3-11,15 leu2-3,112* | MRG7337.1 | “” | 6F |
| MSB203 | *MATa hmr-git1Δ::RS-URA3f-loxP-256xlacO-RS ade2-1::(HIS3p-GFP-lacI::ADE2), cdc20::MET3p-cdc20::loxP MCD1-TAP::natMX leu2-3,112:(GAL1p-RecR::LEU2), trp1-1::(ADH1p-OsTIR1(codon optimized)::TRP1), ura3Δ::AgTEFp-SpHIS5 bar1Δ can1-100 his3-11,15* | MRG7270.1 | “” | 6G |
| MSB204 | *MATa hmr-git1Δ::RS-URA3f-loxP-256xlacO-RS ade2-1::(HIS3p-GFP-lacI::ADE2), cdc20::MET3p-cdc20::loxP MCD1-TAP::natMX leu2-3,112:(GAL1p-RecR::LEU2), trp1-1::(ADH1p-OsTIR1(codon optimized)::TRP1), eco1-3xmAID-5xflag::bleMX ura3Δ::AgTEFp-SpHIS5 bar1Δ can1-100 his3-11,15* | MRG7301.1 | “” | 6G |
| MSB205 | *MATa hmr-git1Δ::RS-URA3f-loxP-256xlacO-RS ade2-1::(HIS3p-GFP-lacI::ADE2), cdc20::MET3p-cdc20::loxP MCD1-TAP::natMX leu2-3,112:(GAL1p-RecR::LEU2), trp1-1::(ADH1p-OsTIR1(codon optimized)::TRP1), wpl1-3xmAID-5xflag::bleMX ura3Δ::AgTEFp-SpHIS5 bar1Δ can1-100 his3-11,15* | MRG7327.1 | “” | 6G |
| MSB206 | *MATa hmr-git1Δ::RS-URA3f-loxP-256xlacO-RS ade2-1::(HIS3p-GFP-lacI::ADE2), cdc20::MET3p-cdc20::loxP MCD1-TAP::natMX leu2-3,112:(GAL1p-RecR::LEU2), trp1-1::(ADH1p-OsTIR1(codon optimized)::TRP1), wpl1-3xmAID-5xflag::bleMX eco1-3xmAID-5xflag::kanMX ura3Δ::AgTEFp-SpHIS5 bar1Δ can1-100 his3-11,15* | MRG7332.1 | “” | 6G |
| MRG7372 | *MATa hmr-git1Δ::RS ade2-1::(HIS3p-GFP-lacI::ADE2), leu2-3,112::(GAL1p-RecR::LEU2), cdc20::MET3p-cdc20::loxP can1-100 his3-11,15 trp1-1 ura3-1* | MSB35.2 | “” | S1B |
| MRG7373 | *MATa hmr-git1Δ::RS ade2-1::HIS3p-GFP-lacI-3xmCherry::hphMX::ADE2::HIS3p-GFP-lacI::ADE2 leu2-3,112::(GAL1p-RecR::LEU2), cdc20::MET3p-cdc20::loxP can1-100 his3-11,15 trp1-1 ura3-1* | MSB167 | “” | S1B |
| MRG7374 | *MATa hmr-git1Δ::RS ade2-1::HIS3p-GFP-lacI-4xmCherry::kanMX::ADE2::HIS3p-GFP-* *lacI::ADE2 leu2-3,112::(GAL1p-RecR::LEU2), cdc20::MET3p-cdc20::loxP MCD1-TAP::natMX ura3Δ::AgTEFp-SpHIS5 can1-100 his3-11,15 trp1-1* | MRG7060 | “” | S1B |
| MRG7320 | *MATa hmr-git1Δ::RS-URA3f-loxP-256xlacO-RS ade2-1::HIS3p-GFP-lacI-3xyeGFP(A206R)::kanMX::ADE2::HIS3p-GFP-lacI::ADE2 leu2-3,112::(GAL1p-RecR::LEU2), cdc20::MET3p-cdc20::loxP MCD1-TAP::natMX ura3Δ::AgTEFp-SpHIS5 can1-100 his3-11,15 trp1-1* | MSB165.1 | “” | S3 |
| MRG7312 | *MATa hmr-git1D::RS-URA3f-lacOPs-RS ade2-1::(HIS3p-GFP-lacI::ADE2), cdc20::MET3p-cdc20::loxP ho::GAL1p-mcd1(R180D,R268D)-TAP::natMX::ho mcd1-3xmAID-5xflag::bleMX ura3Δ::AgTEFp-SpHIS5 bar1Δ can1-100 his3-11,15 trp1-1 leu2-3,112 trp1-1* | MRG6989 | “” | S5A |
| MRG7355.1 | *MATa hmr-git1Δ::RS-URA3f-loxP-256xlacO-RS ade2-1::(HIS3p-GFP-lacI::ADE2), cdc20::MET3p-cdc20::loxP MCD1-TAP::natMX trp1-1::(ADH1p-OsTIR1(codon optimized)::TRP1), wpl1Δ::kanMX eco1-3xmAID-5xflag::bleMX ura3Δ::AgTEFp-SpHIS5 bar1Δ can1-100 his3-11,15 leu2-3,112* | MRG7301.1 | “” | S5A |
| MRG7329 (isolate #1) | *MATa hmr-git1Δ::RS-URA3f-loxP-256xlacO-RS ade2-1::HIS3p-GFP-lacI-3xmCherry::hphMX::ADE2::HIS3p-GFP-lacI::ADE2 cdc20::MET3p-cdc20::loxP MCD1-TAP::natMX trp1-1::(ADH1p-OsTIR1(codon optimized)::TRP1), wpl1-3xmAID-5xflag::bleMX ura3Δ::AgTEFp-SpHIS5 bar1Δ can1-100 his3-11,15 leu2-3,112* | MRG7269.1 |  | S5B |
| MRG6830.1 | *MATa hmr-git1Δ::RS-URA3f-loxP-256xlacO-RS ade2-1::HIS3p-GFP-lacI-3xmCherry::hphMX::ADE2::HIS3p-GFP-lacI::ADE2 leu2-3,112::(GAL1p-RecR::LEU2), cdc20::MET3p-cdc20::loxP MCD1-TAP::natMX ura3Δ::AgTEFp-SpHIS5 wpl1Δ::TRP1 can1-100 his3-11,15 trp1-1* | MSB165.1 | “” | S5B |
| MRG7235.1 | *MATa hmr-git1Δ::RS-URA3f-loxP-256xlacO-RS ade2-1::(HIS3p-GFP-lacI::ADE2), leu2-3,112::GAL1p-SIC1(4m)-AID\*-9xmyc::LEU2 cdc20::MET3p-cdc20::loxP MCD1-TAP::natMX wpl1Δ::TRP1 ura3Δ::AgTEFp-SpHIS5 bar1Δ can1-100 his3-11,15 trp1-1* | MRG7027 | “” | S6A, S6B |
| MRG7232.1 | *MATa hmr-git1Δ::RS-URA3f-loxP-256xlacO-RS ade2-1::HIS3p-GFP-lacI-3xmCherry::hphMX::ADE2::HIS3p-GFP-* *lacI::ADE2 leu2-3,112::GAL1p-SIC1(4m)-AID\*-9xmyc::LEU2 cdc20::MET3p-cdc20::loxP MCD1-TAP::natMX wpl1Δ::TRP1 ura3Δ::AgTEFp-SpHIS5 bar1Δ can1-100 his3-11,15 trp1-1* | MRG7026 | “” | S6A |
| MRG7238.1 | *MATa hmr-git1Δ::RS-URA3f-loxP-256xlacO-RS ade2-1::HIS3p-GFP-lacI-4xmCherry::kanMX::ADE2::HIS3p-GFP-lacI::ADE2 leu2-3,112::GAL1p-SIC1(4m)-AID\*-9xmyc::LEU2 cdc20::MET3p-cdc20::loxP MCD1-TAP::natMX wpl1Δ::TRP1 ura3Δ::AgTEFp-SpHIS5 bar1Δ can1-100 his3-11,15 trp1-1* | MRG7059 | “” | S6A |
| MRG7115 | *MATa hmr-git1Δ::RS-URA3f-loxP-256xlacO-RS ade2-1::HIS3p-GFP-lacI-3xmCherry::hphMX::ADE2 leu2-3,112::GAL1p-SIC1(4m)-AID\*-9xmyc::LEU2 cdc20::MET3p-cdc20::loxP MCD1-TAP::natMX ura3Δ::AgTEFp-SpHIS5 bar1Δ can1-100 his3-11,15 trp1-1* | MRG6919<br>x<br>MRG7041 | “” | S7 |
| MRG7122 | *MATa hmr-git1Δ::RS-URA3f-loxP-256xlacO-RS ade2-1::HIS3p-GFP-lacI-4xmCherry::kanMX::ADE2 leu2-3,112::GAL1p-SIC1(4m)-AID\*-9xmyc::LEU2 cdc20::MET3p-cdc20::loxP MCD1-TAP::natMX ura3Δ::AgTEFp-SpHIS5 bar1Δ can1-100 his3-11,15 trp1-1* | MRG7102<br>x<br>MRG7041 | “” | S7 |

**Table S2.**
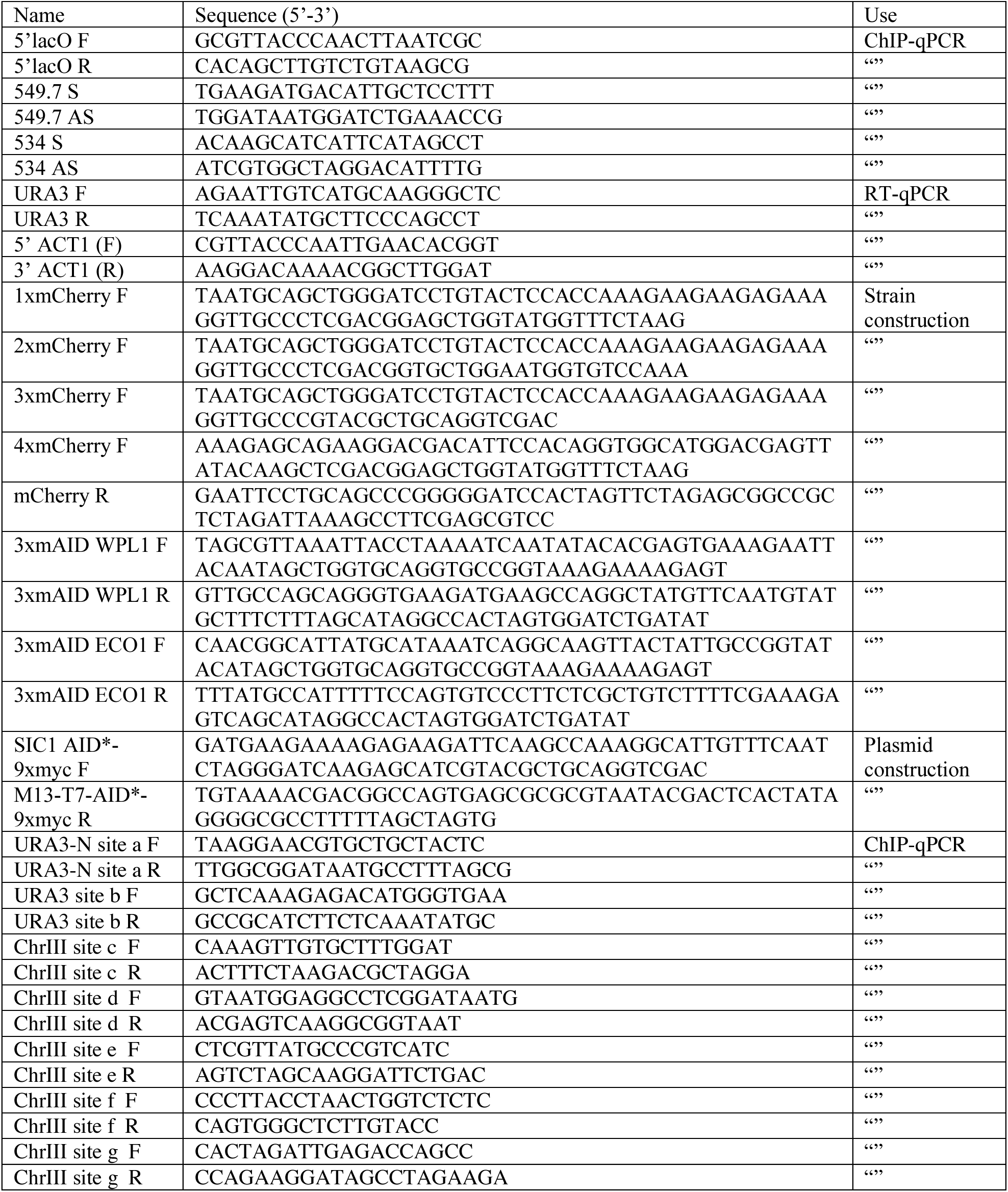
Oligonucleotides.

